# Genolator: A Multimodal Large Language Model Fusing Natural Language, Genomic, and Structural Tokens for Protein Function Interpretation

**DOI:** 10.1101/2025.11.14.688396

**Authors:** Martin Danner, Tanhim Islam, Matthias Begemann, Florian Kraft, Miriam Elbracht, Ingo Kurth, Jeremias Krause

## Abstract

**Background:** Decoding the genetic code to unveil its genome functionality is a monumental task which would greatly advance the understanding of disease mechanisms and development of targeted treatment approaches. Although large language models (LLMs) have transformed natural language processing across diverse domains, translating the complex language of DNA into human-readable form remains challenging due to genomic data complexity and unexplored regions of the human genome. Current (genomic) language models either have a solid understanding of natural language or of the genomic code. Models fusing both aspects are largely lacking.

**Results:** Here we present Genolator, a multimodal large language model that integrates embeddings from DNA sequences, amino acid sequences, and protein structures with natural language queries. Fine-tuned on over 370,000 question-answer pairs generated using abstracted Gene-Ontology (GO) terms, Genolator effectively answers queries regarding protein subcellular localization, molecular function, and biological processes. Evaluation demonstrates high accuracy in confirming or denying protein function associations, outperforming baseline models such as openly available allrounder LLMs like GPT 4.1 as well as smaller domain specific models integrating knowledge from foundation models like Evo2 and ESM-2. Explorations of the Genolator’s hidden states unveil a biologically and linguistically plausible organization of its learned representations.

**Conclusion:** Genolator enhances accessibility to genomic information by enabling natural language interaction with protein data, facilitating biological discovery, and clinical research. It represents a step towards bridging genomic code and human language through the integration of a multimodal LLM.

## 1. Background

Proteins and nucleic acids, alongside lipids and glycans, are fundamental molecules in cells^1,2^. They enable biological functions, store genetic information, and serve as essential building blocks of the cell. The human genome and proteome are vastly complex, and to this day a large portion of the life’s code remains unexplored^2–5^. Yet, deciphering the molecular function of proteins and their involvement in biological processes is key to understanding disease mechanisms and designing targeted treatments^6–8^.

While experimental characterization remains the gold standard for determining protein function, it is inherently resource- and time-intensive, limiting its scalability. To address these constraints, numerous computational methods have emerged to reduce experimental workload and broaden our understanding of the human genome. By prioritizing likely functional candidates for follow-up, these approaches can significantly accelerate downstream experimental validation^7–16^. Among computational approaches, machine learning models are increasingly employed for predicting the functionality and cellular localization of proteins^15^. Part of them are large language models (LLMs) which recently demonstrated significant impact across virtually all domains of life^17^, with applications spanning medicine^18^, education^19^, traffic management^20^, finance^21^ and beyond. Leveraging billions of parameters, these machine learning models can discern intricate patterns in data, enabling them to understand and generate human language and solve a diverse range of complex natural language processing tasks, making them prime candidates to decode the information stored in the genetic code. Therefore, genomic language models have been developed which extend the concept of large language models to genomic language. By adopting LLM-like architectures and incorporating specific adaptations, as in models such as Evo2^22,23^, genomic language models are trained on trillions of DNA base pairs from a wide range of prokaryotic and eukaryotic species^22–27^. This allows them to interpret and understand genetic information, predict the functional impacts of genetic variation, and identify biological features such as exon-intron boundaries, binding sites, and even aspects of protein functionality and structure^22,23^. The knowledge acquired during training is encoded within the model’s parameters, which serve as a foundation for various downstream tasks. However, current genomic language models are mostly inaccessible to non-technical users, need to be finetuned on each downstream task and most importantly lack an understanding of natural language which could prove crucial to decoding the genomic code more easily^28^. Formulating questions about genomic aspects such as genome functionality in natural language and thereby understanding how the information about the genome functionality is encoded in the genetic material, could open a new way to explore and decipher genome biology.

Building on this conceptual foundation and the aforementioned challenges, our study introduces Genolator: an accessible model capable of processing genomic information and natural language in tandem, facilitating the exploration of the human genome through natural language queries. For a proof of concept, we selected the decoding of the functionality of protein coding genes and subsequently proteins as our prime task. Our focus is to enable users to ask and obtain answers to questions about the biological process, molecular function, and cellular component of a given (potentially novel) protein, as derived from the Gene Ontology nomenclature^29^, a standardized framework for the description of protein functionality. In previous work^30^, we demonstrated that sequence embeddings capture significant information but may not encompass all relevant biological features needed for various downstream tasks. To address this, we adopt an intermediate fusion strategy^31,32^, combining four distinct modalities: the DNA sequence, the amino acid sequence, the 3D protein structure of the target gene or protein, and the user’s natural language query.

## 2. Results

### Genolator – a multimodal large language model regarding genome functionality

Genolator is a finetuned multimodal Llama^33–35^ model, designed to fuse natural language with genomic language. It receives a question, formulated in natural language (English), and information from DNA-sequences, amino acid sequences and protein structures in tandem. Each modality is presented to Genolator in the form of an embedding, a latent representation of the encoded information, from a machine learning model trained on the specific modality. DNA sequences are handled by the genomic foundation model Evo2^22^, amino acids by the protein language model ESM-2^36^ and protein structure embeddings created by a graph autoencoder trained on protein structures^30^. Genolator utilizes token projectors to fuse the input from the modality encoders with the natural language question and presents the output to a Llama model which processes the input and formulates a response which will be returned in English. Genolator was designed to answer questions regarding genome functionality. To achieve this, genome functionality descriptions were derived from the three Gene Ontology (GO)-Term^29^ aspects:

- Cellular Component
- Molecular Function
- Biological Processes

which were transformed into a generally understandable phrase by a GPT 4.1 model^37^. Genolator was subsequently trained on over 370,000 question–answer pairs encompassing both genuine protein function associations and denial questions, where incorrect associations were deliberately constructed. The training set included both confirmative and denial questions, which required responses affirming or denying specific associations, as well as open-ended questions demanding more complex, explanatory answers. As a result, Genolator can address both open-ended queries (see Figure 1) and confirmatory or denial questions (see Figure 2) relating to genome functionality.

**Figure 1:**
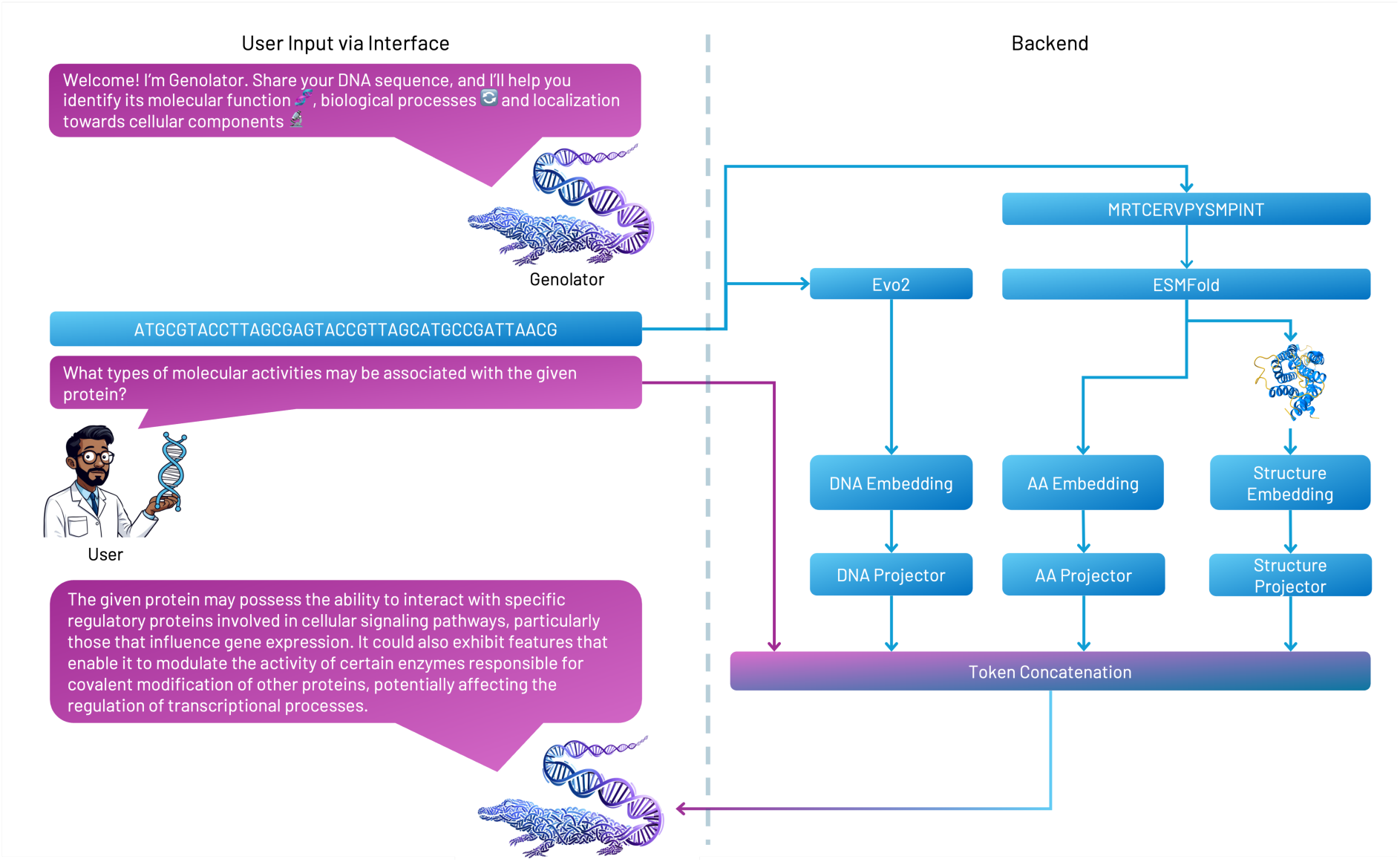
Exemplary illustration of a user assessing the molecular function of a DNA sequence with a generic query. Left: The conversation between Genolator and the user. Right: A schematic overview of the backend process handling the user’s query and parsing of the provided DNA sequence.

**Figure 2:**
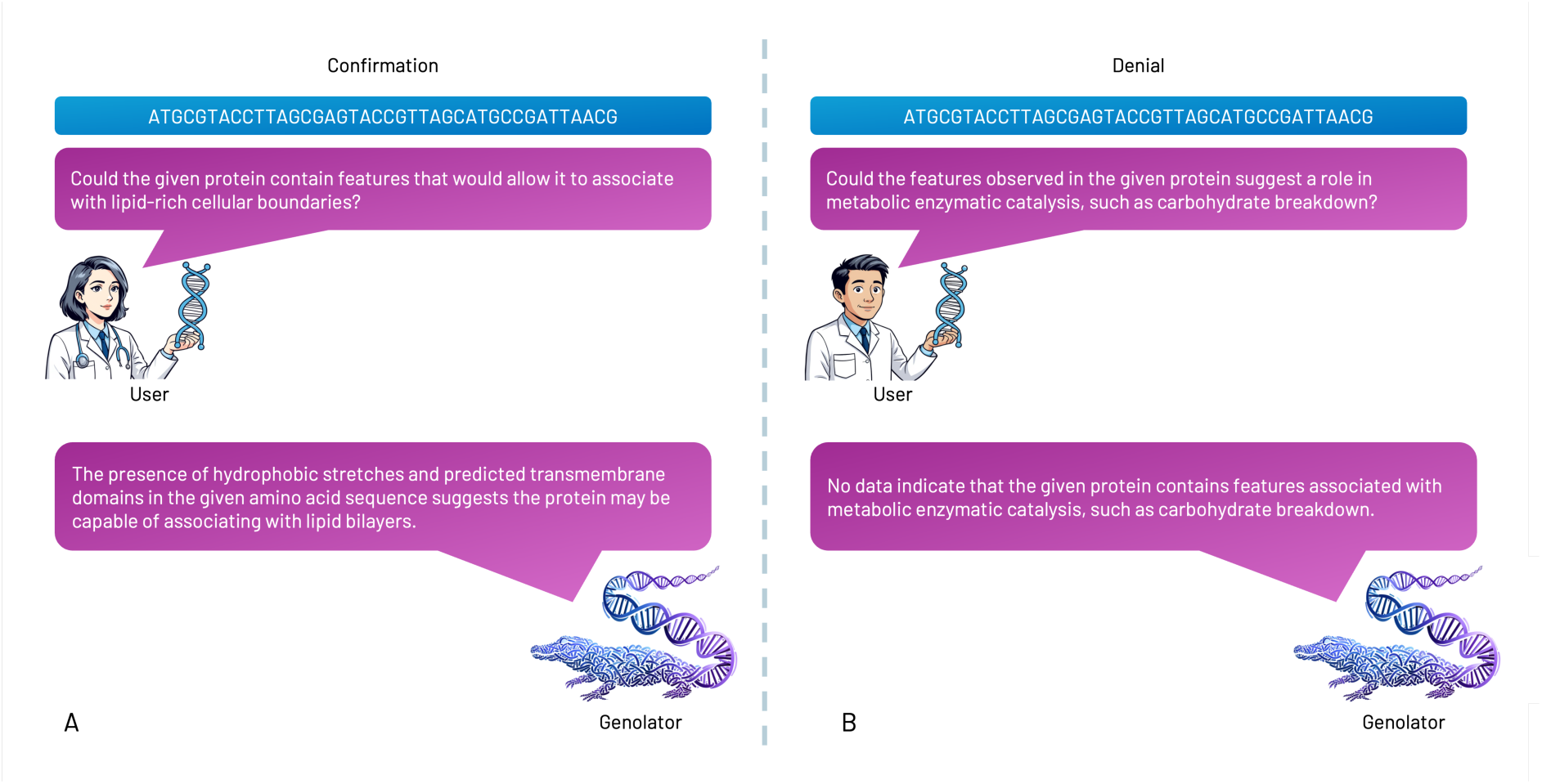
A. Illustration of a sample user interaction with Genolator, in which the user queries whether a sequence of interest is associated with a specific cellular component, and Genolator confirms the association. B. Schematic illustration of a user querying Genolator about the involvement of the given sequence in a particular biological process, which is denied by Genolator.

### Genolator can accurately confirm and deny genome functionality associations and outperforms allrounder and specialist baseline models

To assess the accuracy and validity of Genolator, we first evaluated its performance on a holdout test set comprising 34,743 questions. This set included confirmative and denial questions addressing subcellular localization, molecular function, and involvement in biological processes, specifically for proteins and genes that were entirely unseen during training. Additionally, to evaluate Genolator on proteins with unknown function, we employed a custom similarity-aware KMeans-based^38^ data split. This splitting strategy ensured that test set samples were not only unseen during training but also resided in distinct regions of the joint multimodal embedding space, thus minimizing structural or sequence similarity to the training data across all modalities. To evaluate Genolator’s performance on the task to answer denial and confirmative questions the task was reframed as a binary classification problem. The performance was evaluated using Accuracy, Precision, Recall, F1-Score and the Matthews-Correlation-Coefficient (MCC)^39^. Genolator (full), in which both the virtual token projectors and the Llama model were jointly trained/fine-tuned, achieved an overall F1-Score of around 0.977. For both the confirmation and denial of associated terms, it reached a MCC of 0.955, the full results are displayed in Table 1. For further quantification of Genolator’s performance, multiple baseline models were benchmarked on the test set. These included general-purpose “all-rounder” LLMs such as GPT-4.1, locally deployable models like the 8B version of Llama-3^33^, and a smaller, domain-specific XGBoost^40^ classifier trained on information from foundation models (Evo2, ESM-2) together with BioBERT^41^ sentence-transformer embeddings of the input questions. Additionally, we evaluated an ablated version of Genolator (TP), in which only the virtual token projectors were trained while the Llama^33^ backbone remained frozen. For GPT 4.1^37^ and the non-finetuned Llama-3^33^ the DNA sequence and the amino acid sequence was directly embedded into the prompt. Genolator outperformed all baseline models. GPT 4.1^37^ reached a MCC of 0.18 and the XGBoost model reached a MCC of 0.933. The ablated version of Genolator (TP) reached a MCC of 0.869. The non finetuned Llama model had to be excluded from the performance comparison. Due to the O(n²) complexity of the self-attention mechanism, a non-multimodal Llama^33^ model encounters memory limitations and excessive computational time when processing large, unembedded DNA sequences. Even when deployed on a high-memory GPU such as the NVIDIA H100, the model faces significant issues with long input sequences, making practical inference infeasible for these inputs. For cases where the base model was capable to produce an answer, the answer highlighted further difficulties of the base model, these being:

1. False remembrance
2. Amino-Acid gibberish
3. Answering the question with a follow up question.

**Table 1:** Performance of Genolator (and Baseline Models) on Confirmation/Denial Questions

|  | Precision | Recall | F1-Score | Support | Model |
| --- | --- | --- | --- | --- | --- |
| Confirmation | 0.984351 | 0.970903 | 0.977581 | 15225.00 | Genolator (full) |
| Denial | 0.970660 | 0.984219 | 0.977392 | 14891.00 | Genolator (full) |
| Accuracy | 0.977487 | 0.977487 | 0.977487 | 0.977487 | Genolator (full) |
| Macro Avg. | 0.977506 | 0.977561 | 0.977487 | 30116.00 | Genolator (full) |
| Weighted Avg. | 0.977582 | 0.977487 | 0.977488 | 30116.00 | Genolator (full) |
| MCC | 0.955066 | 0.955066 | 0.955066 | 0.955066 | Genolator (full) |
| Confirmation | 0.585473 | 0.650115 | 0.616103 | 15225.00 | GPT 4.1<sup>37</sup> |
| Denial | 0.596775 | 0.529412 | 0.561079 | 14892.00 | GPT 4.1<sup>37</sup> |
| Accuracy | 0.590431 | 0.590431 | 0.590431 | 0.590431 | GPT 4.1<sup>37</sup> |
| Macro Avg. | 0.591124 | 0.589763 | 0.588591 | 30117.00 | GPT 4.1<sup>37</sup> |
| Weighted Avg. | 0.591062 | 0.590431 | 0.588895 | 30117.00 | GPT 4.1<sup>37</sup> |
| MCC | 0.180882 | 0.180882 | 0.180882 | 0.180882 | GPT 4.1<sup>37</sup> |
| Confirmation | 0.967463 | 0.962463 | 0.964957 | 15478.00 | XGBoost<sup>40</sup> |
| Denial | 0.966166 | 0.970688 | 0.968422 | 17092.00 | XGBoost<sup>40</sup> |
| Accuracy | 0.966779 | 0.966779 | 0.966779 | 0.966779 | XGBoost<sup>40</sup> |
| Macro Avg. | 0.966815 | 0.966575 | 0.966689 | 32570.00 | XGBoost<sup>40</sup> |
| Weighted Avg. | 0.966782 | 0.966779 | 0.966775 | 32570.00 | XGBoost<sup>40</sup> |
| **MCC** | 0.933389 | 0.933389 | 0.933389 | 0.933389 | XGBoost<sup>40</sup> |
| **Confirmation** | 0.906075 | 0.968359 | 0.936182 | 15202.00 | Genolator (TP) |
| **Denial** | 0.965261 | 0.897522 | 0.930160 | 14891.00 | Genolator (TP) |
| **Accuracy** | 0.933307 | 0.933307 | 0.933307 | 0.933307 | Genolator (TP) |
| **Macro Avg.** | 0.935668 | 0.932941 | 0.933171 | 30093.00 | Genolator (TP) |
| **Weighted Avg.** | 0.935362 | 0.933307 | 0.933202 | 30093.00 | Genolator (TP) |
| **MCC** | 0.868604 | 0.868604 | 0.868604 | 0.868604 | Genolator (TP) |

In case one it seemed like the base model tried to utilize knowledge from its initial training, which was ultimately futile and lead to associations with wrong proteins. In case two the base model ignored the question and returned a generated DNA or Amino Acid Sequence, in a next character-prediction style. In some cases, after generating some characters the model fell into a repeating pattern of amino acids. In case three the base model also ignored the question and returned a (unrelated) question.

Because predicting genome functionality varies in difficulty across the three Gene Ontology (GO) aspects - molecular function, cellular component, and biological process- we evaluated and compared Genolator’s (full) performance with the baseline models for each category. For all models, associating a protein with a molecular function proved to be the easiest task, yielding the highest MCCs. Except for GPT-4.1^37^, which performed better on the biological process aspect, the cellular component task ranked second in difficulty, while predicting involvement in biological processes was the most challenging aspect overall. While the ablated projector-only version of Genolator (TP) model outperformed GPT-4.1^37^, it was surpassed by the smaller XGBoost^40^ model.

### The non-finetuned publicly available GPT 4.1 model (falsely) remembers genomic data and struggles with the concept of sequence similarity

Publicly available large language models (e.g., ChatGPT) can, in principle, be applied (and probably are applied) to genomic data. However, our previous results show that general-purpose models such as GPT-4.1^37^ and Llama-3^33^ (8B parameters) underperform compared to modality-specific approaches like XGBoost^40^ or Genolator. To further investigate this limitation, we conducted a focused case study analysing the ability of GPT-4.1^37^ to process nucleotide and amino acid sequences when these are directly embedded within the prompt. For this we presented GPT 4.1^37^ with nucleotide sequences and amino acid sequences of well characterized genes such as BRCA1 (as a representative for proteins with more than 1000 amino acids), IHH (as a representative for proteins with an intermediate length) and INS, encoding insulin, (as a representative for small proteins) and asked the model openly formulated questions about the involvement of the corresponding protein in biological processes, the associated molecular functions and the cellular localisation of the protein. Since GPT 4.1^37^ was trained on large volumes of publicly available data, it is possible (and likely) that the model already encountered the genomic information during training. Therefore, in addition to the genomic functionality questions, we also prompted the model to identify the provided nucleotide and amino acid sequence, resulting in three identification queries per case. The summarized results are presented in Table 2. For the INS gene, the GPT-4.1^37^ model was able to correctly identify the provided DNA and amino acid sequences. Additionally, it accurately described all assigned molecular functions, biological process involvements, and the correct subcellular localization for INS. For longer sequences such as BRCA1 and IHH, GPT 4.1^37^ claimed that it performed a BLAST^42^ search which led it to the identification of the allegedly correct identification of the BRCA1 sequence as TP53 and HTT, while IHH was identified as COL1A1 and MPZ. As shown in Table 2, the proteins identified by GPT-4.1^37^ for BRCA1 and IHH do not exhibit significant sequence similarity to the reference sequences. This suggests that GPT-4.1^37^ may have difficulty interpreting sequence similarity in context, leading to the misidentification of these proteins. Next to issues with sequence similarities, the subsequent molecular functions were also partly incorrect, as BRCA1 was describes as a protein involved in transport across membranes.

**Table 2:**
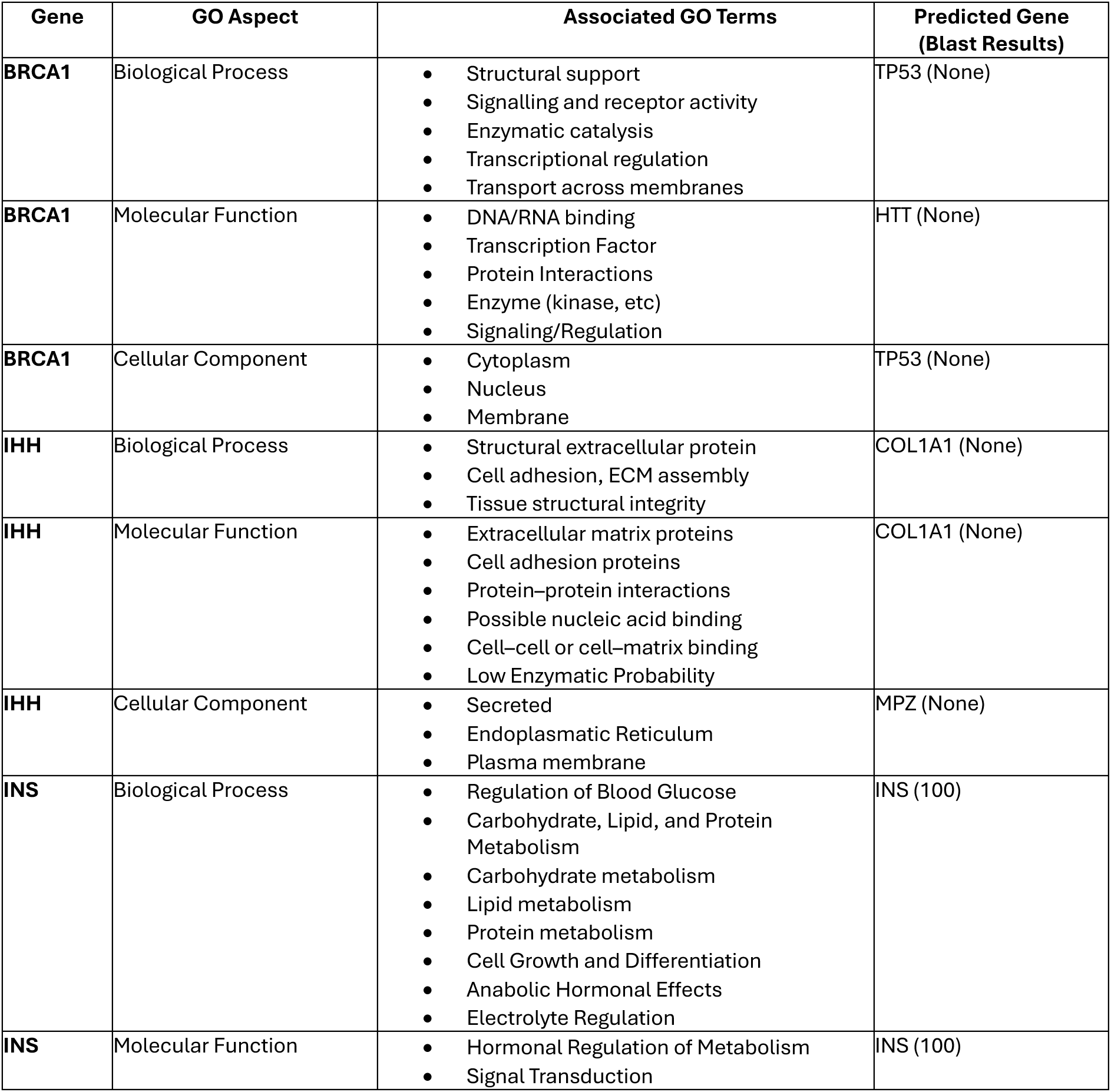
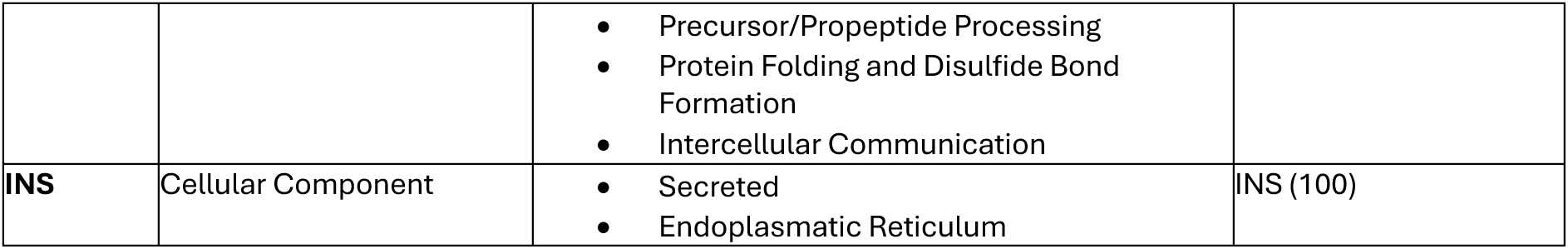
Results of the small case study of GPT 4.1 capabilities to analyse nucleotide sequences and amino acid sequences for aspects of genome functionality.

| Gene | GO Aspect | Associated GO Terms | Predicted Gene (Blast Results) |
| --- | --- | --- | --- |
| **BRCA1** | Biological Process | • Structural support<br>• Signalling and receptor activity<br>• Enzymatic catalysis<br>• Transcriptional regulation<br>• Transport across membranes | TP53 (None) |
| **BRCA1** | Molecular Function | • DNA/RNA binding<br>• Transcription Factor<br>• Protein Interactions<br>• Enzyme (kinase, etc)<br>• Signaling/Regulation | HTT (None) |
| **BRCA1** | Cellular Component | • Cytoplasm<br>• Nucleus<br>• Membrane | TP53 (None) |
| **IHH** | Biological Process | • Structural extracellular protein<br>• Cell adhesion, ECM assembly<br>• Tissue structural integrity | COL1A1 (None) |
| **IHH** | Molecular Function | • Extracellular matrix proteins<br>• Cell adhesion proteins<br>• Protein–protein interactions<br>• Possible nucleic acid binding<br>• Cell–cell or cell–matrix binding<br>• Low Enzymatic Probability | COL1A1 (None) |
| **IHH** | Cellular Component | • Secreted<br>• Endoplasmatic Reticulum<br>• Plasma membrane | MPZ (None) |
| **INS** | Biological Process | • Regulation of Blood Glucose<br>• Carbohydrate, Lipid, and Protein Metabolism<br>• Carbohydrate metabolism<br>• Lipid metabolism<br>• Protein metabolism<br>• Cell Growth and Differentiation<br>• Anabolic Hormonal Effects<br>• Electrolyte Regulation | INS (100) |
| **INS** | Molecular Function | • Hormonal Regulation of Metabolism<br>• Signal Transduction | INS (100) |
|  |  | • Precursor/Propeptide Processing<br>• Protein Folding and Disulfide Bond Formation<br>• Intercellular Communication |  |
| **INS** | Cellular Component | • Secreted<br>• Endoplasmatic Reticulum | INS (100) |

### Genolator’s hidden states contain linguistically and biological plausible representations

To assess whether the model effectively integrated natural language and biological information, we investigated the features learned by Genolator. We extracted the hidden states from the final layer of the Llama^33^ model prior to the output layer and projected them into a two-dimensional space using t-distributed stochastic neighbour embedding (t-SNE)^43^. This analysis was performed separately for each of the three GO-term^29^ aspects, allowing us to visualize the distribution of hidden states according to whether the model confirmed or denied the queried association. Excerpts from the resulting visualizations can be seen in the figures 3-5. For most GO terms, Genolator achieved a clear separation between confirmed and denied associations in the t-SNE^43^ space, producing distinct clusters across all GO-term aspects^29^. Additionally, within each GO aspect^29^, Genolator clustered related terms in close proximity, while clearly separating less related terms. For example, as visible in figure 3, for the association with biological processes the term “regulation of DNA-templated transcription” is directly neighboured by the term “chromatin organization”, a necessary preceding step for DNA translation, and the term “cytoplasmic translation”, a procedure following transcription. Cell cycle and cell death related processes were also clustered closely, neighboured by the term “DNA repair”. Similarly, developmental related biological processes were grouped together. Additionally, signalling and immune related biological processes were clustered accordingly. For the presented subset of the category molecular function, a similar observation was made, as visible in figure 4. In the selected subset visible in figure 4, we observed clustering between DNA and RNA related functionalities (“DNA binding”, “RNA binding”, “histone binding” and “transcription regulator activity”), structure related functionalities (“protein folding chaperones” and “structural molecule activity”), signalling and transport related functionalities (“transporter activity”, “receptor ligand activity” and “transducer activity”) and enzymes (“ligase activity”, “hydrolase activity” and “transferase activity”). Finally, for cellular components we again observed a plausible structuring of the hidden states, as visible in figure 5. In this subset, nuclear compartment related localisations (“chromosome”, “nuclear chromosome”, “nucleolus”, “nucleus” and “nucleoplasm”) were grouped together, similarly to extracellular compartments (“extracellular matrix”, “extracellular region”, “extracellular space”).

**Figure 3:**
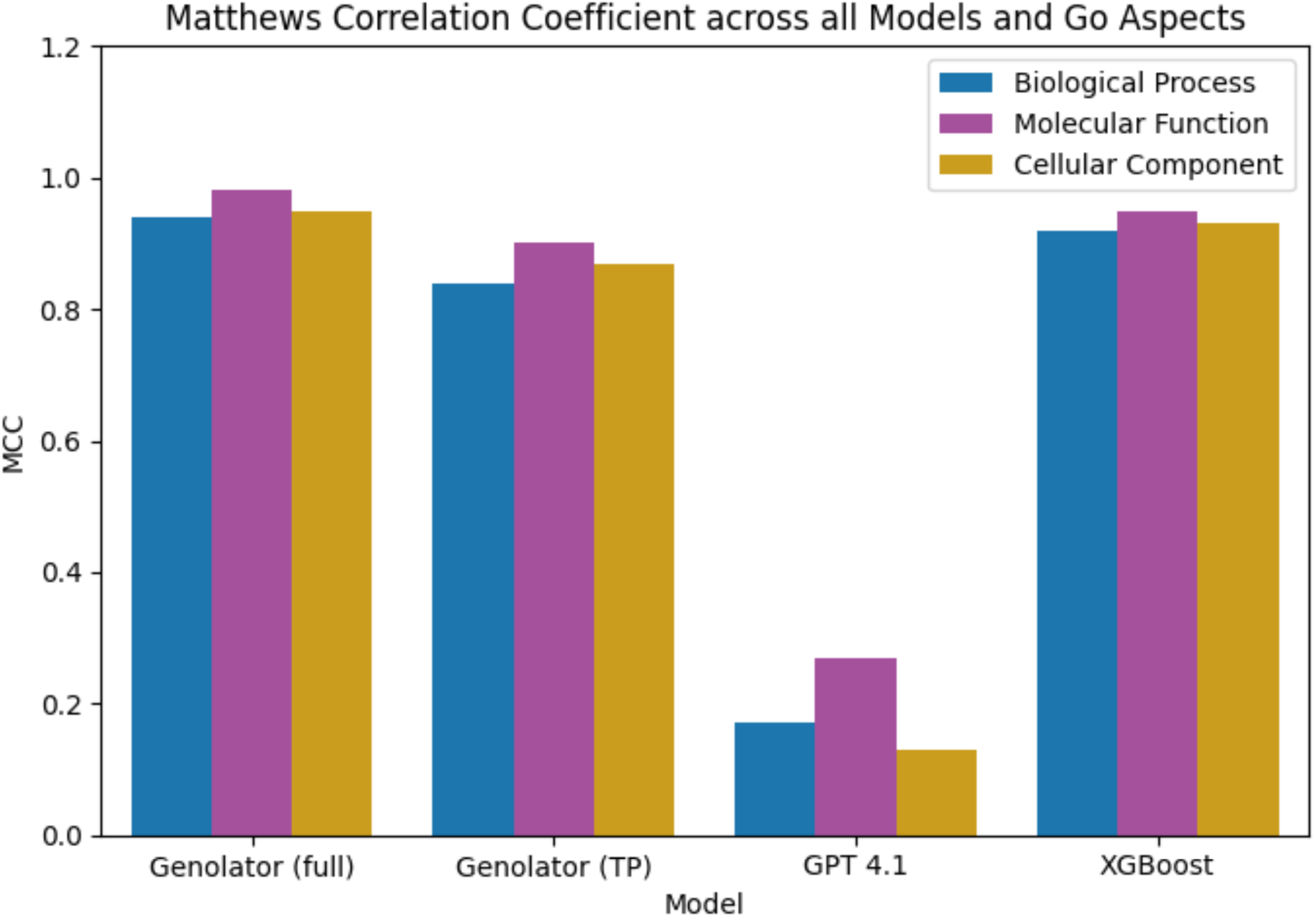
Bar plot displaying the Matthews Correlation CoePicient (MCC) achieved by each model across diPerent Gene Ontology (GO) Aspects. The x-axis denotes the various models evaluated, while the y-axis indicates the MCC values. Each colored bar represents a diPerent GO Aspect: Blue – Biological Process, Purple – Molecular Function, Yellow – Cellular Component

**Figure 3:**
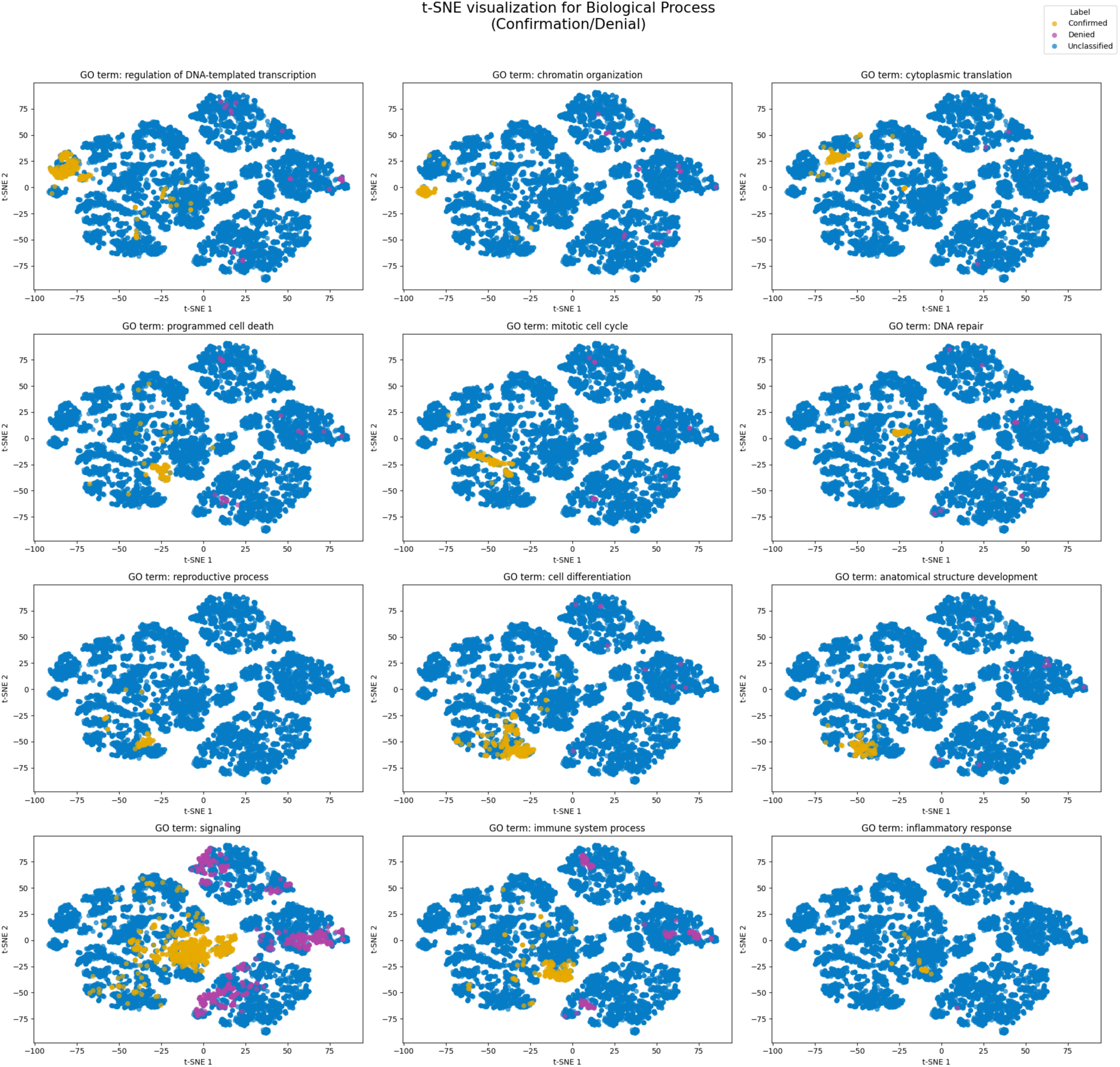
Figure 3: t-SNE visualization for a selection of Biological Process Gene Ontology (GO) terms^29^, generated using the last hidden states from the fine-tuned Llama^33^ model. Each subplot represents a diPerent GO term^29^, displaying embeddings projected into two dimensions (t-SNE 1 and t-SNE 2 axes). Data points are colored as follows: yellow (“Confirmed”) indicates the model associates the corresponding gene with the GO term^29^; purple (“Denied”) indicates the model denies the association; blue (“Unclassified”) indicates genes for which no association was queried.

**Figure 4:**
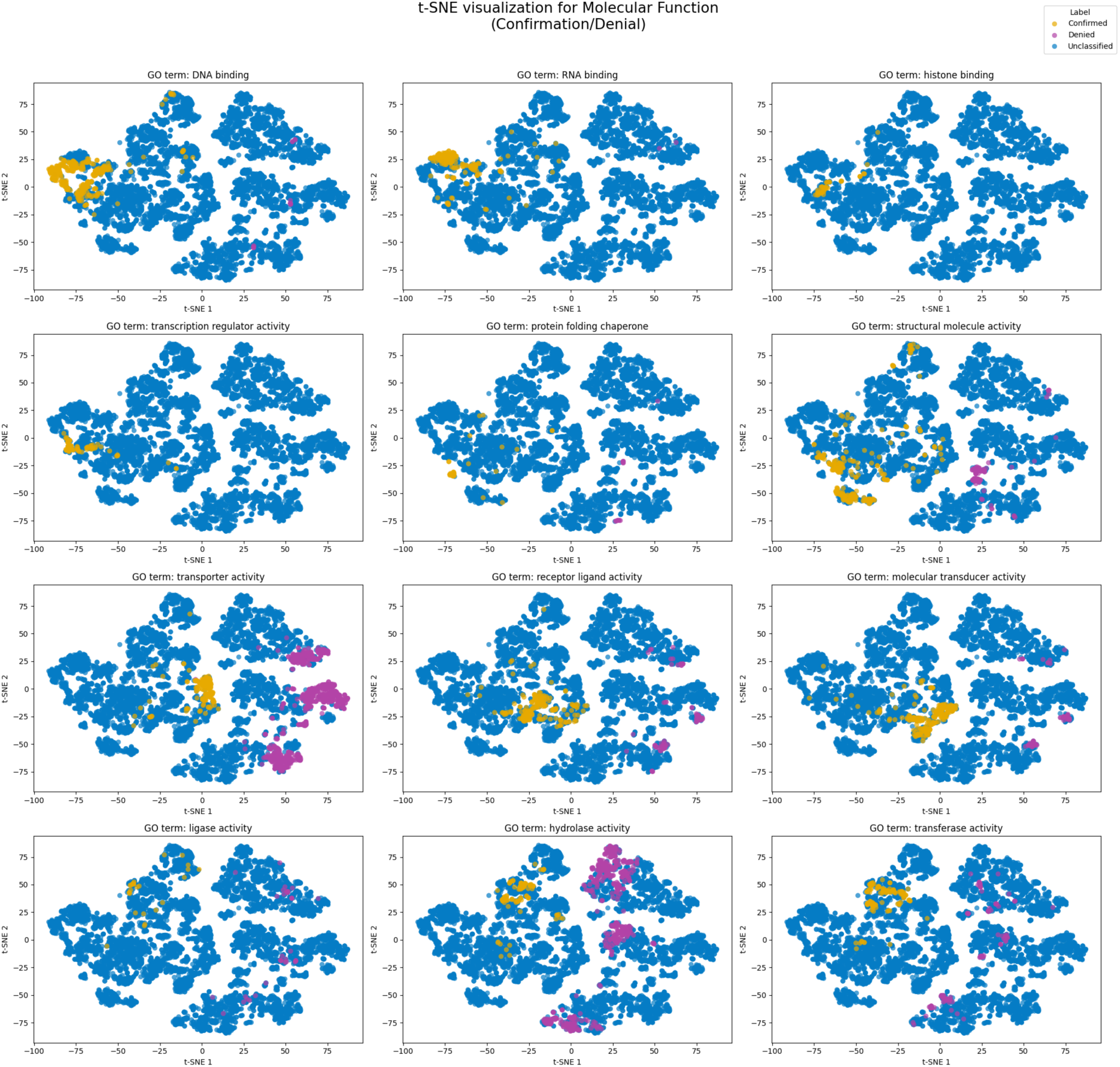
t-SNE visualization for a selection of Molecular Function Gene Ontology (GO) terms^29^, generated using the last hidden states from the fine-tuned Llama^33^ model. Each subplot represents a diPerent GO term^29^, displaying embeddings projected into two dimensions (t-SNE 1 and t-SNE 2 axes). Data points are colored as follows: yellow (“Confirmed”) indicates the model associates the corresponding gene with the GO term^29^; purple (“Denied”) indicates the model denies the association; blue (“Unclassified”) indicates genes for which no association was queried.

**Figure 5:**
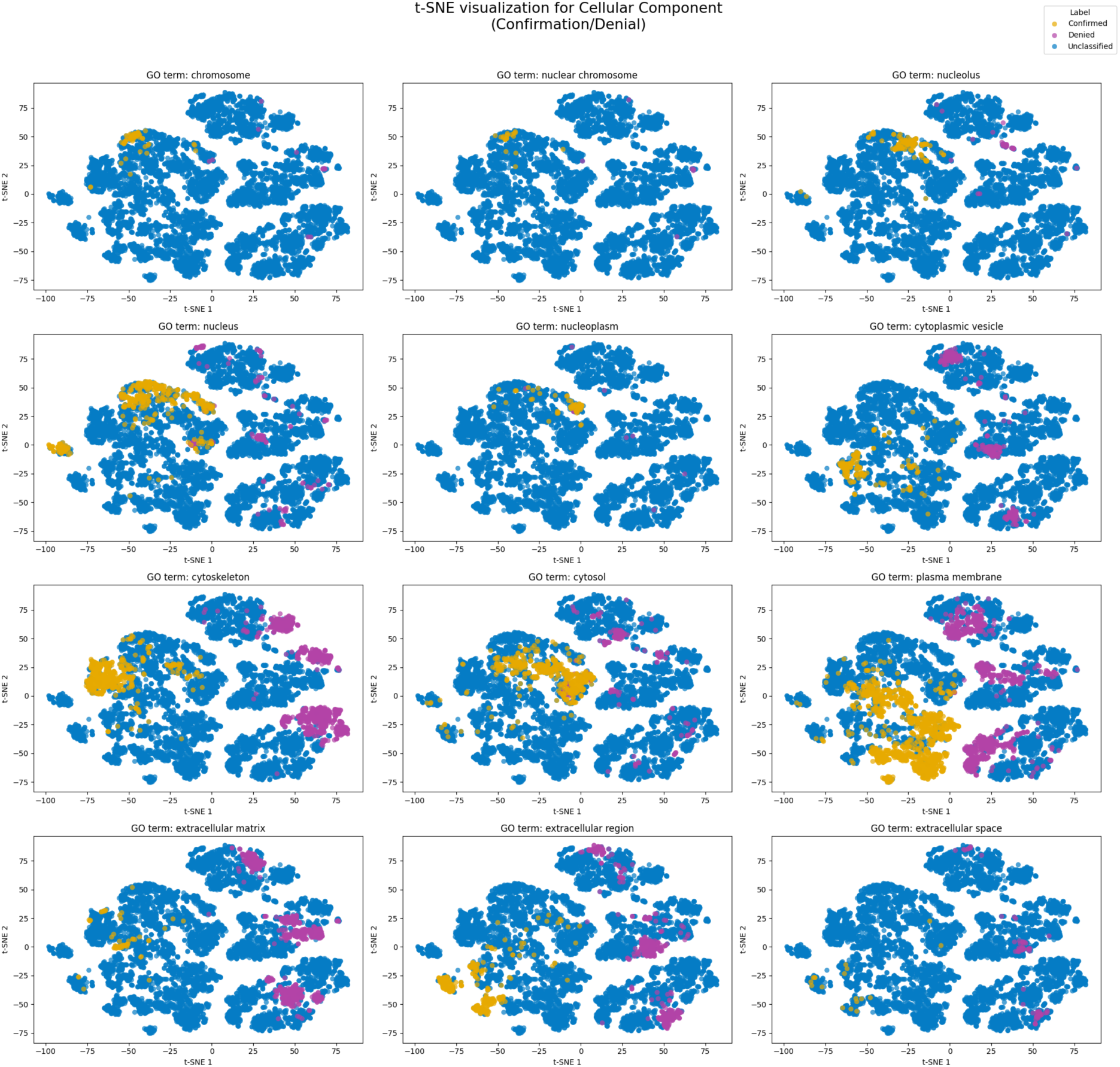
t-SNE visualization for a selection of Cellular Component Gene Ontology (GO) terms^29^, generated using the last hidden states from the fine-tuned Llama^33^ model. Each subplot represents a diPerent GO term^29^, displaying embeddings projected into two dimensions (t-SNE 1 and t-SNE 2 axes). Data points are colored as follows: yellow (“Confirmed”) indicates the model associates the corresponding gene with the GO term^29^; purple (“Denied”) indicates the model denies the association; blue (“Unclassified”) indicates genes for which no association was queried.

**Figure 6:**
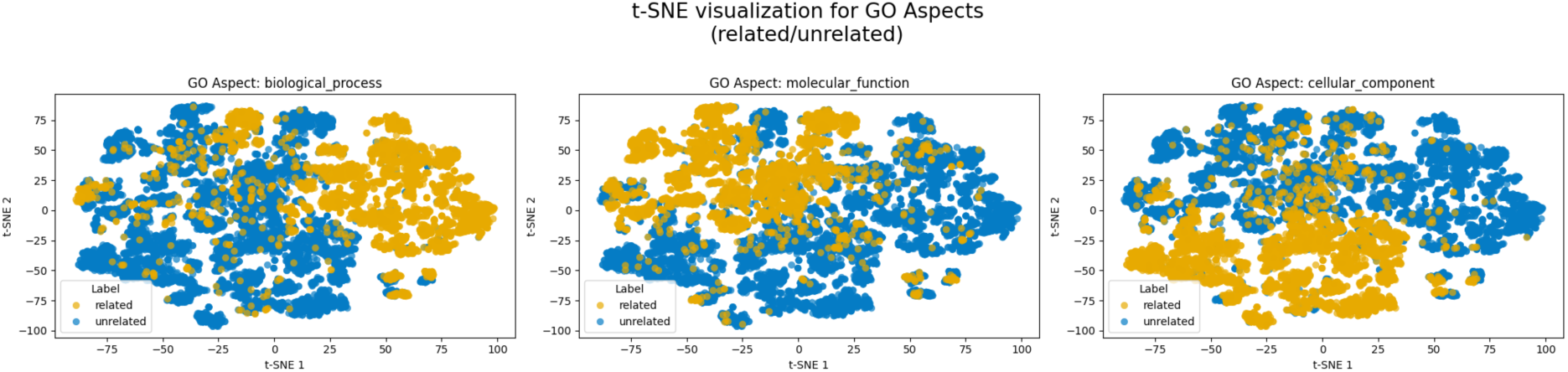
t-SNE^43^ visualizations of the confirmation questions, generated from the last hidden states of the fine-tuned Llama model. Each subplot corresponds to one of the three GO aspects^29^. Yellow points represent datapoints related to relevant (confirmed) questions for the respective GO aspect^29^, while blue points denote unrelated datapoints.

### The Genolator implicitly learns the diEerent categories of the GO-hierarchy

To further investigate Genolator’s general language and genomic understanding, we analysed its hidden states to assess whether the model could distinguish GO terms^29^ belonging to different aspects. As shown in Figure 7, Genolator was predominantly able to form three largely distinct clusters, each corresponding to one of the three GO aspects^29^.

**Figure 7:**
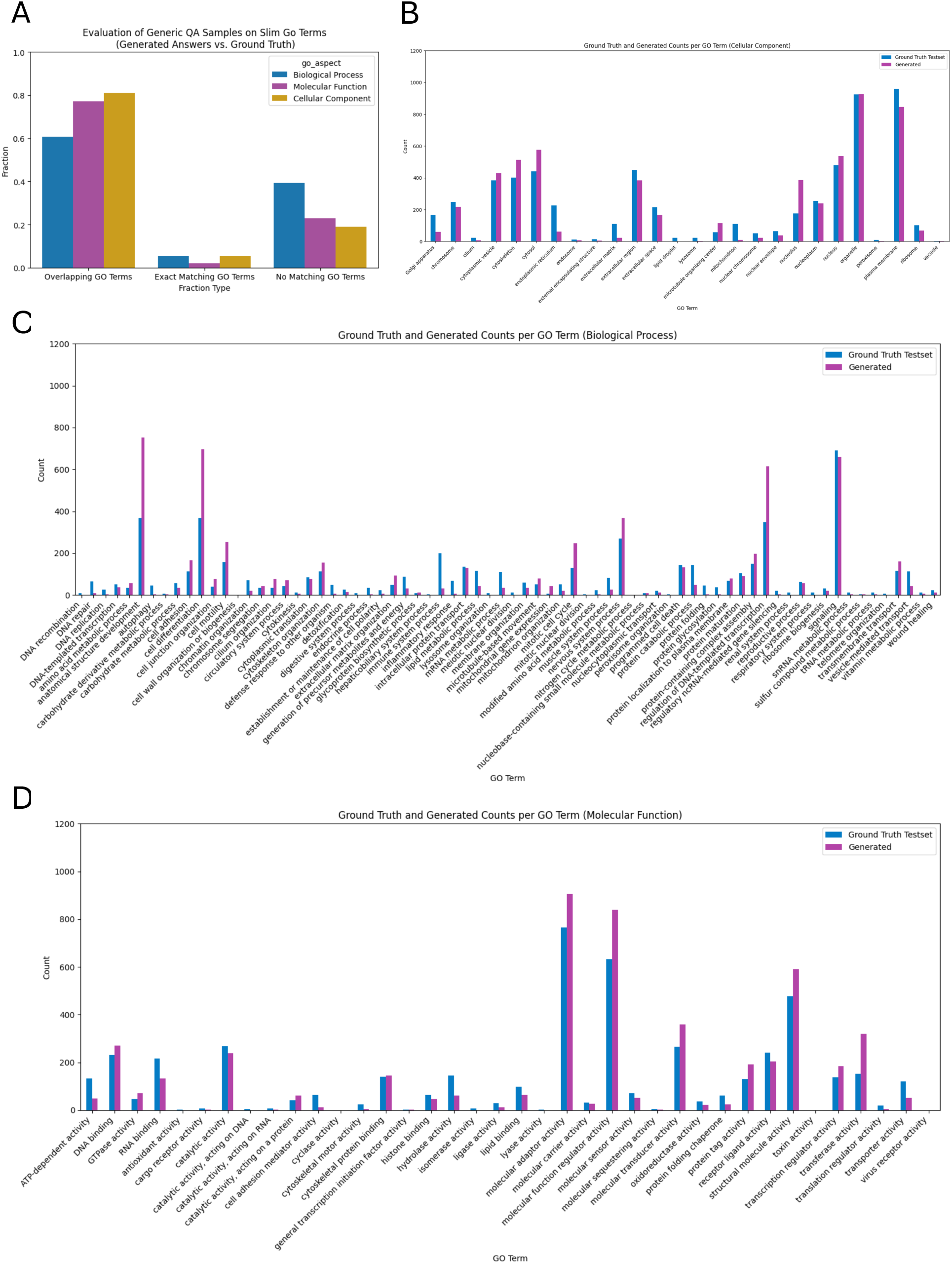
A) Bar plot illustrating the evaluation of open question-answer (QA) samples on slim Gene Ontology (GO) terms^29^, comparing generated answers to ground truth annotations. The x-axis shows diPerent fraction types: Overlapping GO Terms^29^, Exact Matching GO Terms^29^, and No Matching GO Term^29^. B) Bar plot showing the counts of ground truth and generated answers for each GO term^29^ within the Cellular Component aspect. C) Bar plot showing the counts of ground truth and generated answers for each GO term^29^ within the Biological Process aspect. D) Bar plot showing the counts of ground truth and generated answers for each GO term^29^ within the Molecular Function aspect. For panels B–D, each GO^29^ term is displayed on the x-axis, with the corresponding sample counts on the y-axis (ranging from 0 to 1200). For each GO term^29^, bars are grouped and color-coded: blue represents the Ground Truth counts and purple represents the Generated answer counts.

### Asking the Genolator openly formulated questions about genome functionality

Next to understanding the confirmation or denial of associations between GO aspects^29^ and specific proteins, we also trained Genolator to answer open-ended questions about GO aspects^29^. This open-ended, more generic capability is particularly valuable for proteins whose involvement in certain GO aspects is currently unclear or poorly characterized (“uncharacterized proteins”). For this we created a separate QA-dataset. We mapped all GO aspects to the slim GO terms using a GPT 4.1^29^ model, thereby enabling the evaluation of the overlap between the predicted and ground truth values. Here we differentiated between three groups, depending on the correctness of the prediction made by Genolator compared to the ground truth. The first group included samples with an exact match, where all ground truth functional terms were correctly predicted, and no additional terms were introduced. The second group comprised samples with partial overlap - some, but not all, ground truth terms were predicted, or additional, irrelevant terms were included. The final group contained samples in which only terms absent from the ground truth were predicted. The visualisation for the three GO aspects^29^ can be seen in figure 8A. For generic questions regarding the involvement in biological processes 5.39 % perfect matches were achieved, 60.78 % matches showed an overlap with the ground truth and 39.22 % showed a complete dissonance between the prediction and the ground truth. For the prediction of the cellular component 5.32 % perfect matches were achieved, 80.94 % showed an overlap with the ground truth and 19.06 % were in complete disagreement with the ground truth. Finally, for the prediction of the molecular function, 2.14 % of the samples showed a perfect match, 77.15 % showed an overlap with the ground truth and 22.85 % were in complete disagreement.

**Figure 8:**
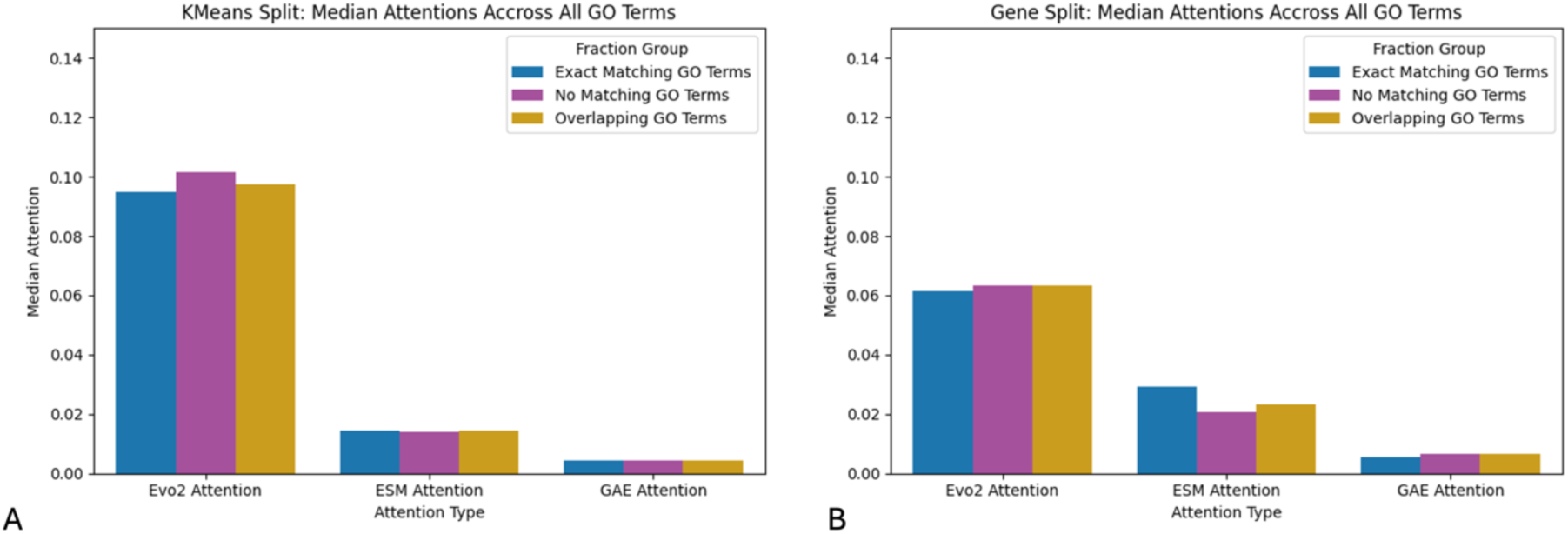
A) Depicting the median attention score, across all GO-terms^29^, separated by GO-aspects^29^ for the KMeans^38^ split. B) Depicting the median attention scores, across all GO-terms^29^, separated by GO-aspects^29^ for the gene name split.

In the Cellular Component aspect (Figure 8B), both the ground truth and Genolator’s predicted answers prominently feature terms such as “organelle” and “plasma membrane” each occurring more than 800 times. In contrast, less common terms like “lipid droplet”, “lysosome”, “peroxisome”, and “endosome” appear fewer than 50 times. Notably, the overall distribution of GO Slim term^29^ counts is similar between ground truth and Genolator outputs, emphasizing the model’s effectiveness in recapitulating the relative frequencies of cellular components.

In the Biological Process aspect (Figure 8C), both the ground truth and Genolator’s predicted answers most prominently feature the term “signalling”, which appears more than 600 times in each. Other frequently occurring terms, such as “anatomical structure development”, “cell differentiation”, and “regulation of DNA-templated transcription”, occur fewer than 350 times in the ground truth, but are predicted over 700 times by Genolator, highlighting a notable disparity. Terms like “DNA recombination”, “meiotic nuclear division”, and “tRNA metabolic process” are rare in both datasets, each with fewer than 50 occurrences. Overall, the distribution of GO Slim term^29^ counts in the biological process category is less well matched between predicted and ground truth answers than in the cellular component aspect.

In the Molecular Function aspect (Figure 8D), “molecular adaptor activity” is the most frequently predicted term by Genolator, with more than 700 occurrences substantially exceeding its count in the ground truth. The second most frequent term, “molecular function regulator activity” also appears more than 600 times in Genolator’s predictions, again outnumbering its ground truth representation. In contrast, terms such as “antioxidant activity”, “cyclase activity” and “virus acceptor activity” are among the least frequent, appearing only rarely in both predicted and ground truth datasets. Overall, the agreement between ground truth and Genolator’s predictions for molecular function is better than for biological processes and is nearly as close as observed in the cellular component aspect.

### Analysing Attention distribution across open questions

To assess the potential benefits of combining Evo2^22^, ESM-2^36^, and protein autoencoder embeddings^30^, we evaluated how multimodal integration influenced Genolator’s predictions for open-ended questions. Specifically, we analysed the model’s attention scores across the different GO aspects. Overall, Evo2^22^ embeddings consistently received the highest attention, followed by ESM-2^36^ embeddings, while protein structure embeddings contributed the least among the three modalities. This visualization is presented in Figure 9. To assess the impact of dataset splitting strategy on model attention distribution, we repeated the analysis using a model trained with a gene-wise data split instead of the previously established KMeans^38^ approach. Under this gene-based split, we observed a noticeable decrease in attention toward the Evo2^22^ embeddings and a corresponding increase in attention directed toward the ESM-2^36^ embeddings.

**Figure 9:**
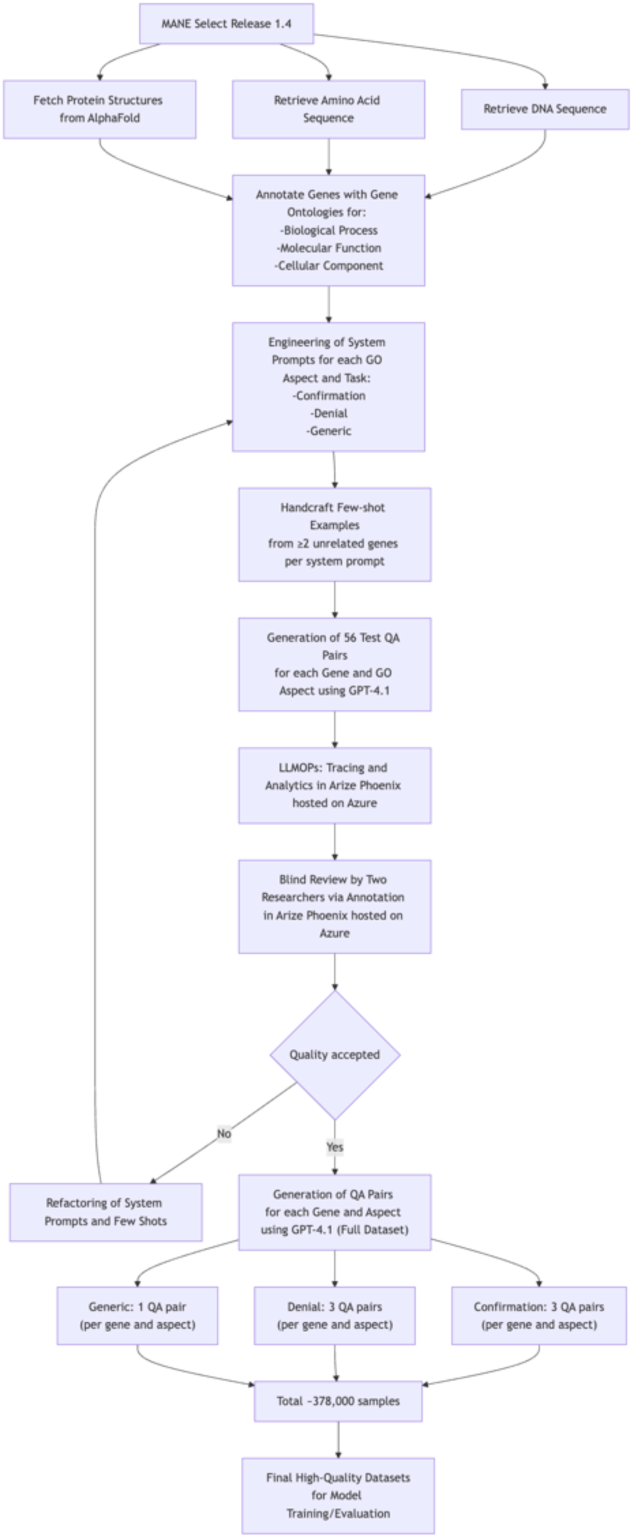
QA Dataset creation workflow

**Figure 10:**
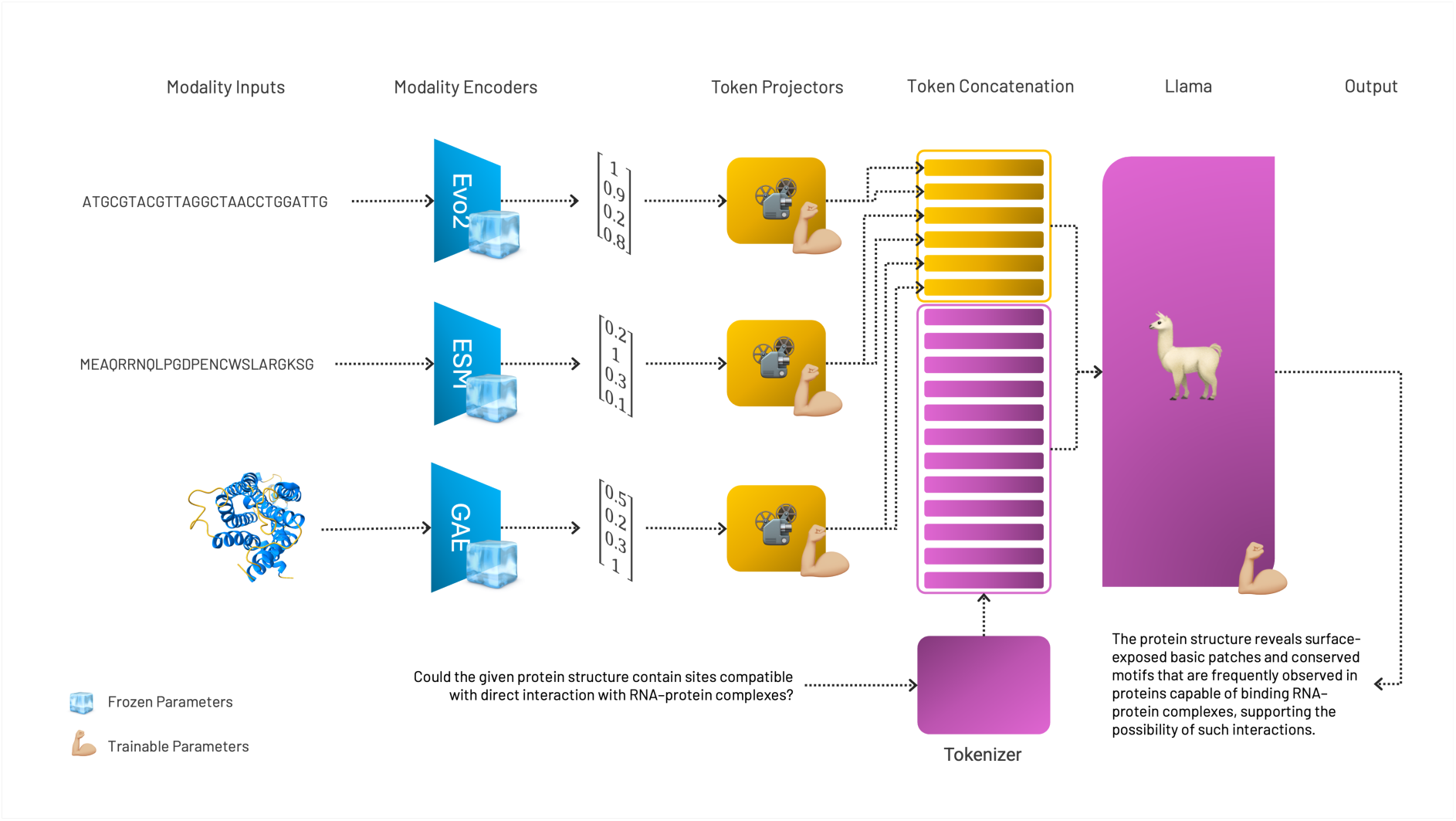
Schematic Illustration of the multimodal fine tuning using feature fusion via token concatenation with tokens derived from genomic as well as protein sequences and protein structures.

## 3. Discussion

### The Genolator – a multimodal language model - has learned the vocabulary of genome biology

Here we present the Genolator, a finetuned Llama^33^ model capable of giving robust evaluations whether a specific protein is associated with functionality terms derived from the three GO aspects^29^ cellular localisation, molecular function and involvement in biological processes. We demonstrate that integrating a large language model capable of processing natural language with genomic embeddings from multimodal models yields a system that outperforms both general-purpose LLMs such as GPT-4.1^37^ and smaller, domain-specific models such as an XGBoost^40^ classifier trained on the same embeddings. The resulting Genolator can interpret genomic embeddings based on instructions given in English, thereby making the use of embeddings from large genomic models more accessible and paving the way for a new, language-driven interface for interacting with genomic data. Furthermore, by fusing natural language with genomic code, Genolator is able to learn interpretable representations that cluster related GO terms^29^ in a biologically meaningful manner. This plausibility is maintained not only for linguistically similar terms but also for biologically related processes and compartments, highlighting the potential of natural language as a powerful tool for exploring genome function. Previously it has been demonstrated that the prediction of GO-terms^29^ can be improved by including explainable text descriptions to the predicting model^7,9^. The development and evaluation of Genolator build upon this evidence, as its strong performance on confirmative and denial questions, as well as the analysis of its hidden states in these contexts, provide additional support for the value of integrating language-based explanations in functional genomics prediction. Consistent with previous reports, we found that predicting the association between a genomic element and a biological process was the most challenging aspect of genome functionality prediction^7^.

However, the difference to the other two GO-categories was small. Kulmanov et al. reported that accurate classification of a protein’s involvement in biological processes requires that the predictive model learn which other proteins are present in the organism’s proteome^7^. This suggests that Genolator was able to leverage such contextual information from its training data. Further studies are required to transfer the Genolators applicability to multiple species and observe how this ability is influenced by proteins from multiple species being present in the training data.

### The Genolator outperforms generalist and specialist baseline models

We provide extensive evidence that openly available LLMs such as GPT-4.1^37^ are currently inadequate for processing genomic sequences and truly understanding the underlying genomic context; their capabilities are limited to natural language-based comprehension. On top of this, these models often generate responses with extensive explanations and may claim to use genomic analysis tools such as BLAST^42^ to lend credibility to their answers, even when their outputs are incorrect or contain hallucinated elements, including fabricated BLAST^42^ results. Additionally, with its large context length (1.000.000 tokens), it should have the technical capability to process large genomic sequences. However, in our small case study we observed that GPT 4.1^37^ only handles small proteins well (such INS), for which it might remembered the complete amino acid sequence. For longer proteins such as BRCA1, it does not only retrieve dissimilar proteins (as indicated by the non-significant overlap returned by BLAST^42^) but also suggest candidates which are dissimilar in the simplest aspects such as the amino acid length (BRCA1 has around 1800 amino acids, HTT has around 3000 amino acids). Combining all this evidence with its poor performance on the binary classification task (MCC = 0.18) suggests that GPT 4.1^37^ struggles with the analysis of genomic sequences. These findings are particularly intriguing given that GPT-4.1^37^ has likely encountered a substantial number of genomic sequences during its training. This observation underscores that – as already stated - despite such exposure, its capabilities remain limited to natural language-based comprehension rather than a true understanding of genomic context. This highlights the need for multimodal language models with a more nuanced grasp of genomic information such as Genolator.

In addition to GPT-4.1, we compared Genolator to an XGBoost^40^ model, which was provided with the same genomic embeddings as well as prompt embeddings from a BioBERT^41^ sentence transformer. Although Genolator outperformed the XGBoost^40^ model across all GO-term categories, the advantage of Genolator extends well beyond this modest performance margin. By leveraging a generative model like Llama, Genolator enables natural language querying, making the tool more accessible and facilitating the use of natural language to explore its learned representations. It should also be noted that, for cost reasons, the Llama-based Genolator was not subjected to hyperparameter tuning, while the XGBoost^40^ model did undergo tuning, giving the latter a slight advantage in terms of optimization. Tuning Genolator’s hyperparameters and playing with the number of virtual tokens offers significant potential for further performance improvement. Moreover, replacing the Llama^33^ backbone with larger, state-of-the-art language models (e.g. GPT-5^44,45^ or Gemini^46,47^) could also provide substantial benefits.

### DiEerentiating to existing models

The field of integrating genomic language models with natural language models is still emerging. To our knowledge, the only similar model is ChatNT^46^, a generalist conversational agent that leverages the nucleotide transformer model to address various binary and regression tasks involving genomic features and elements. However, ChatNT^28^ has not been trained to predict genomic functionality with respect to GO aspects^29^ and does not provide the capability to incorporate amino acid embeddings or protein structure information. Furthermore, Evo2^22^, the model used to generate DNA embeddings for Genolator, offers a substantially larger input size (1 million) compared to the nucleotide transformer model (12 kb)^24^. This was especially advantageous for Genolator, as it enabled the encoding of complete genes, including introns and UTR regions, which on average exceed the input length limitations of the nucleotide transformer^24^. Both increasing the sequence context window and integrating embeddings for amino acid sequences and protein structure have been identified as important future directions by the creators of ChatNT^28^; however, these features are currently absent from their model. The two approaches also differ fundamentally in their training strategies. In Genolator, only the token projectors and the Llama^33^ model were trained/fine-tuned, while all embedding models - including Evo2^22^, ESM-2^36^, and the protein structure encoder^30^ -remained frozen throughout training. By contrast, ChatNT^28^ trains both its projection model and its nucleotide encoder (analogous to Genolator’s use of Evo2^22^ for the DNA embedding step).

A key advantage of our approach is the demonstrated value of foundation models like Evo2^22^, which can generate highly informative embeddings for downstream tasks without requiring additional fine-tuning. This flexibility allows for rapid migration or replacement of embedding models, as well as straightforward retraining. The strong performance of our XGBoost^40^ baseline, trained directly on these genomic and protein embeddings, further underscores their quality.

Additionally, we evaluated an ablated version of Genolator in which only the token projectors were trained while keeping the Llama^33^ backbone frozen. Although this projector-only model underperformed relative to the fully fine-tuned Genolator and the XGBoost^40^ model, it still achieved remarkably good results suggesting a knowledge distillation-like^48^ effect. This is particularly noteworthy, as it enables efficient migration and experimentation with larger LLMs: only the token projectors need to be retrained, greatly reducing computational costs and making the approach highly efficient.

One limitation of our approach is that neither the architecture nor the hyperparameters of the token projectors (nor those of the Llama model in the case of the full Genolator) were subject to extensive tuning, which suggests further room for optimization in future work. Further studies will be necessary to systematically compare these different strategies and to establish optimal training protocols for multimodal natural language and genomic language models.

Finally, it is important to note that ChatNT^28^ is specifically designed for targeted binary classification tasks (such as detecting the presence of genomic elements like lncRNAs) and regression tasks (such as predicting protein stability), where it demonstrates strong performance. However, it is not intended to answer generic, open-ended natural language questions. This specialized focus allows ChatNT^28^ to excel in well-defined prediction scenarios, while also underscoring the need for further research to explore the capabilities and adaptability of generalist genomic conversational agents in addressing open-ended functional genomics queries.

### Evaluating the Genolators capability to answer open questions

Training a large language model to answer generic open-ended questions, rather than confirming or denying specific functional associations, broadens the range of possible applications. However, it comes with some hurdles. One challenge is to assess the quality of the generated responses. In general, it is possible to evaluate the predictions made by employing experts from the relevant field, this however was not realistically possible for a project like this in which multiple thousands of predictions are made encompassing a wide topic, due to limited personnel. To address these challenges, we evaluated Genolator’s predictions from multiple perspectives to provide a comprehensive overview of its capabilities in answering generic open questions. LLMs are commonly evaluated using cross entropy and perplexity^49^. While perplexity is a standard metric for evaluating language models, it has significant limitations in open-ended tasks such as answering generic questions about biological processes, molecular functions, and cellular components of a given sequence. Perplexity and cross-entropy do not directly capture the quality or correctness of generated responses. This challenge is reflected in recent findings that perplexity, while widely used as a proxy for evaluation, cannot be directly linked to factuality in generation settings, as it is affected by many linguistic phenomena and by the variability of valid responses^50–52^. The issue is particularly pronounced for proteins with a wide range of known functions, as the ground truth dataset contains many associated labels, making it considerably more challenging for the model to identify every single label. Moreover, the specific phrasing of a response can impact both perplexity and cross-entropy, since different textual descriptions of the same function may not be evaluated fairly by these metrics - a limitation observed in several cases produced by Genolator.

To address these shortcomings, we implemented a GPT-4.1-based LLM-as-judge approach to map both Genolator’s predictions and the ground truth annotations to a reduced set of GO Slim terms^29^, thereby reducing evaluation complexity and enhancing comparability. Additionally, when constructing the training samples, we simplified the ground truth GO term descriptions to focus only on the primary GO terms for each gene, facilitating fairer and more consistent assessments. However, this approach may sometimes disadvantage Genolator during evaluation, as it is possible for the model to correctly predict legitimate GO terms for a gene that were not included in the shortened ground truth annotations. Overall, while perplexity remains useful for intra-model comparisons, a more nuanced and factuality-aware evaluation is essential for reliable assessment and continued improvement of model and dataset quality in open-ended biological tasks.

We observed that, for most questions regarding proteins in the test dataset (biological processes: 60.78%; molecular function: 77.15%; cellular localization: 80.94%), Genolator predicted functionalities that overlapped with the ground truth. The count comparisons demonstrate that Genolator effectively captures the relative distribution of GO Slim terms^29^ for cellular component and molecular function, closely mirroring the ground truth labels. However, greater divergence in the biological process aspect highlights remaining challenges in accurately predicting more complex or diverse functional categories. This demonstrates Genolator’s capability to answer open-ended questions, while also indicating areas where further improvement -as well as the development of more robust evaluation strategies - is possible. Importantly, our analysis brings to light a critical challenge in modeling and evaluating protein function: the significant underrepresentation of certain GO terms^29^ in the available data. This data imbalance not only complicates fair evaluation but may also limit the model’s ability to generalize to rare or less frequently annotated functions. An additional benefit of incorporating open questions into Genolator’s training set may be the enhancement of its general language understanding. As described above, Genolator was able to clearly distinguish between questions related to different GO aspects^29^, a skill it may have developed through exposure to open-ended, generic questions during training.

### Attention analysis reveals a limited benefit of multimodality

Genolator is a multimodal LLM that integrates embeddings from various genomic modalities. Across our experiments, we consistently observed a preference for Evo2^22^ embeddings over the others. Several factors may account for this observation. Firstly, the DNA embeddings (Evo2^22^) had the highest information content, as they incorporated non-coding regions (such as introns and UTRs) and were produced by the most extensive of the three foundation models. Notably, Evo2^22^ embeddings had a dimensionality of 4096, compared to 2560 for amino acid sequence embeddings (ESM-2^36^) and just 128 for the structure embeddings.

For model input, each embedding modality was mapped to 8 virtual tokens of size 4096. Consequently, projecting the highly condensed structural embeddings (128 dimensions) into a much larger space (4096 dimensions) may not provide as much meaningful information as the already high-dimensional Evo2^22^ embeddings. This dimensionality mismatch likely contributed to lower attention being assigned to the structure modality.

It is plausible that using a structural encoder capable of producing less condensed, higher-dimensional embeddings could shift the distribution of attention, potentially improving Genolator’s performance. Furthermore, systematically varying the number of virtual tokens according to the modality, rather than using a fixed number, could be another avenue for optimization.

Due to resource constraints, our study was limited to two train-test splits using different splitting criteria. Notably, the relative importance of Evo2^22^ versus ESM-2^36^ embeddings varied substantially between splits, suggesting that attention patterns may depend on sequence content and split methodology. Nevertheless, these findings indicate that incorporating multiple modalities might be beneficial and highlights the value of further explorations of multimodal approaches within the Genolator framework.

Further work is needed to systematically explore the impact of embedding dimensionality, encoder architecture, and virtual token allocation on multimodal integration and the overall performance of genomic language models like Genolator.

## 4. Conclusion

Genolator is a generative machine learning model, fusing natural language with the language of life, the genetic code. We highlight that Genolator can use its language processing capabilities while also getting a solid grasp in processing genomic information, as observable in its strong performance in the confirmation and denial question dataset. An analysis of its hidden states reveals that Genolator has a fused understanding of both languages, as visible in the provided t-SNE^43^ plots.

Importantly, this work underscores the technical achievement in multimodal integration and demonstrates new ways to interact with genomic data. As natural language interfaces continue to advance, tools like Genolator pave the way for more intuitive and accessible engagement with complex biological information expanding the possibilities for research, diagnostics, and discovery, particularly regarding poorly characterized genomic regions.

While Genolator already demonstrates strong performance, the adoption and training of even larger and more powerful language models might yield further improvements. Continued research will focus on enhancing Genolator’s capacity to answer open-ended questions and broadening its applicability to an ever-wider range of genomic and biomedical tasks.

## 5. Methods

### Construction of Custom QA Datasets

As a foundation for our QA datasets, we utilized the MANE (Matched Annotation from NCBI and EMBL-EBI^53^) Select release 1.4 (GRCh38), which provides single representative transcripts for each canonical protein-coding gene. For each gene, we retrieved both the DNA sequence using BioPython^54^ and the corresponding amino acid sequence. Genes were further annotated with gene ontologies (GO)^29^ encompassing all three aspects: Molecular Function, Cellular Component, and Biological Process.

To ensure the integrity of our fine-tuning and evaluation, we explicitly constrained the data generation process: gene names, gene symbols, the raw DNA sequences, and amino acid sequences were never included/provided in any generated QA samples, prompts, or natural language context. This measure was taken to ensure that the model cannot leverage prior learned knowledge about genes by gene names or recognizable DNA/amino acid sequences to infer the correct response, thereby enforcing that all answers are derived from reasoning over the multimodal integration of provided genomic DNA, amino acid and protein structure embedding.

For each GO aspect^29^, we engineered specific system prompts to guide a GPT-4.1^37^ model (hosted on Azure AI Foundry) in generating three confirmation QA samples per gene, leveraging the ontology annotations and at least six manually crafted few-shot examples from two or more unrelated genes per prompt. These samples formed the confirmation QA datasets.

To construct denial QA samples, we applied an analogous strategy: three denial samples were generated per aspect and gene by instructing the GPT-4.1^37^ model to formulate questions regarding associations with GO terms that were not present in the genés ground truth annotations. This approach ensured that denial samples specifically addressed molecular functions, biological processes or cellular components not – at least so far – attributed to the target gene.

Furthermore, for each GO aspect^29^ and gene, we also engineered a generic QA sample using a dedicated system prompt designed to elicit broad, open-ended questions and answers about potential biological processes, molecular functions or cellular components that could be associated with a given sequence or structure. This prompt excludes mention of specific gene names, protein names, pathway names, mechanisms, or nucleic acid terminology, and focuses instead on plausible broad processes/functions/components, with responses tailored to reflect the main biological process, molecular function and cellular component associated with the GO term^29^ provided.

In total, we produced nine datasets:

- Confirmation QA: Biological Process
- Confirmation QA: Cellular Component
- Confirmation QA: Molecular Function
- Denial QA: Biological Process
- Denial QA: Cellular Component
- Denial QA: Molecular Function
- Generic QA: Biological Process
- Generic QA: Cellular Component
- Generic QA: Molecular Function

Each confirmation and denial dataset comprised approximately 54,000 QA pairs (three per ∼18,000 genes), while each generic dataset consisted of 18,000 QA samples (one per gene), yielding a total of around 378,000 samples across all datasets. To ensure and estimate dataset quality, QA samples from a random selection of 56 genes were independently and blindly reviewed by two researchers, who annotated each sample as “accepted,” “declined,” or “review.”

For this and to support LLM evaluation and monitoring, we employed Arize Phoenix, an open-source platform for LLM operations, trace collection, and analytics. A dedicated Phoenix instance was self-hosted on Azure using Azure App Service for deployment, SQLite in combination with Azure Data Lake Gen2. This infrastructure enabled systematic analysis, annotation, and evaluation of the LLM generated QA samples throughout the project lifecycle. Lastly, Protein structural data was obtained from the AlphaFold Protein Structure Database^55–57^ in form of PDB-files.

### Genomic Sequence Embeddings with Evo2

To obtain compact genomic representations, we utilized Evo2^22^ 7B, a large-scale genomic foundation model trained on 9.3 trillion DNA base pairs. For each gene’s DNA sequence, per-token embeddings were extracted from the ‘blocks.28.mlp.l3’ layer, following the model authors’ recommendations. These token-level embeddings were then aggregated using mean pooling along the sequence dimension, resulting in a fixed-size 4096-dimensional embedding vector for each gene. Evo2^22^ was deployed and inference was conducted on an Azure Standard_NC40ads_H100_v5 virtual machine, equipped with 40 CPU cores, 320 GB RAM, and a single NVIDIA H100 GPU.

### Protein Sequence Embeddings with ESM-2

Amino acid sequence embeddings were generated using the ESMFold^36^ (3B) model accessed via HuggingFace^58^ and deployed on an Azure Standard_NC24ads_A100_v4 virtual machine (24 CPU cores, 220 GB RAM, and one NVIDIA A100 GPU). Due to the O(n³) complexity of the ESMFold^36^ model, longer sequences occasionally led to memory overflows. To address this, protein sequences exceeding 1,000 amino acids were divided into subsequences of 1,000 residues each, with model inference performed separately on each segment.

For each token of the sequence, embeddings from the last hidden layer of the ESM-2 block were extracted and concatenated, yielding per-token vectors of length 2,560. These token-level embeddings were aggregated via mean pooling along the sequence dimension, resulting in a single 2,560-dimensional embedding vector per gene.

### Structural Embeddings via Graph Autoencoders

Structural protein embeddings were generated using a Graph Convolutional Autoencoder (GCAE) trained on over 60,000 protein structures predicted by ESMFold^36^, as described by Danner et al.^30^ As it could be demonstrated this model is also compatible with AlphaFold-derived structures. Thus, for each gene, the protein structure of the MANE transcript was retrieved from the AlphaFold Protein Structure Database. The corresponding PDB files were first converted to graph representations using Graphein^59^. These graphs were then processed by the GCAE, assigning a 128-dimensional embedding to each node. Node embeddings were subsequently aggregated by mean pooling, resulting in a single 128-dimensional embedding vector.

### Multimodal Embedding Integration and Llama Fine-Tuning

As a base model we selected Bio-Medical-Llama-3 8B provided by ContactDoctor on Hugging Face, an 8-billion parameter version of Llama-3 that is pre-fine-tuned on biomedical samples for improved natural language understanding in medical and genetics contexts.

### Multimodal Projection into the Llama Embedding Space

To enable multimodal integration within the Llama architecture, we mapped the embeddings representing the DNA sequences (Evo2^22^), amino acid sequences (ESM-2^36^), and protein structures (Graph Convolutional Autoencoder) into Llama’s native token embedding space. This was achieved by designing three separate virtual token projectors, one per modality, with identical architecture. Each projector applied a learnable linear transformation to map the input embedding dimension to the hidden size expected by Llama (4096). For each sample, eight virtual tokens per modality were produced, yielding a total of 24 multimodal virtual tokens, which were concatenated and prepended to the textual input tokens for downstream model processing.

Followingly, all token projectors, as well as the Llama^33^ model itself, were fine-tuned jointly on the QA datasets. During training, the weights of the original foundation models (used to generate the modality-specific embeddings) remained frozen, ensuring that only the virtual token projectors and the Llama model were updated. Figure 11 provides a visual overview of the proposed multimodal model architecture, including the virtual token projection and integration of DNA, sequence, and structure embeddings within the Llama backbone.

**Figure 11:**
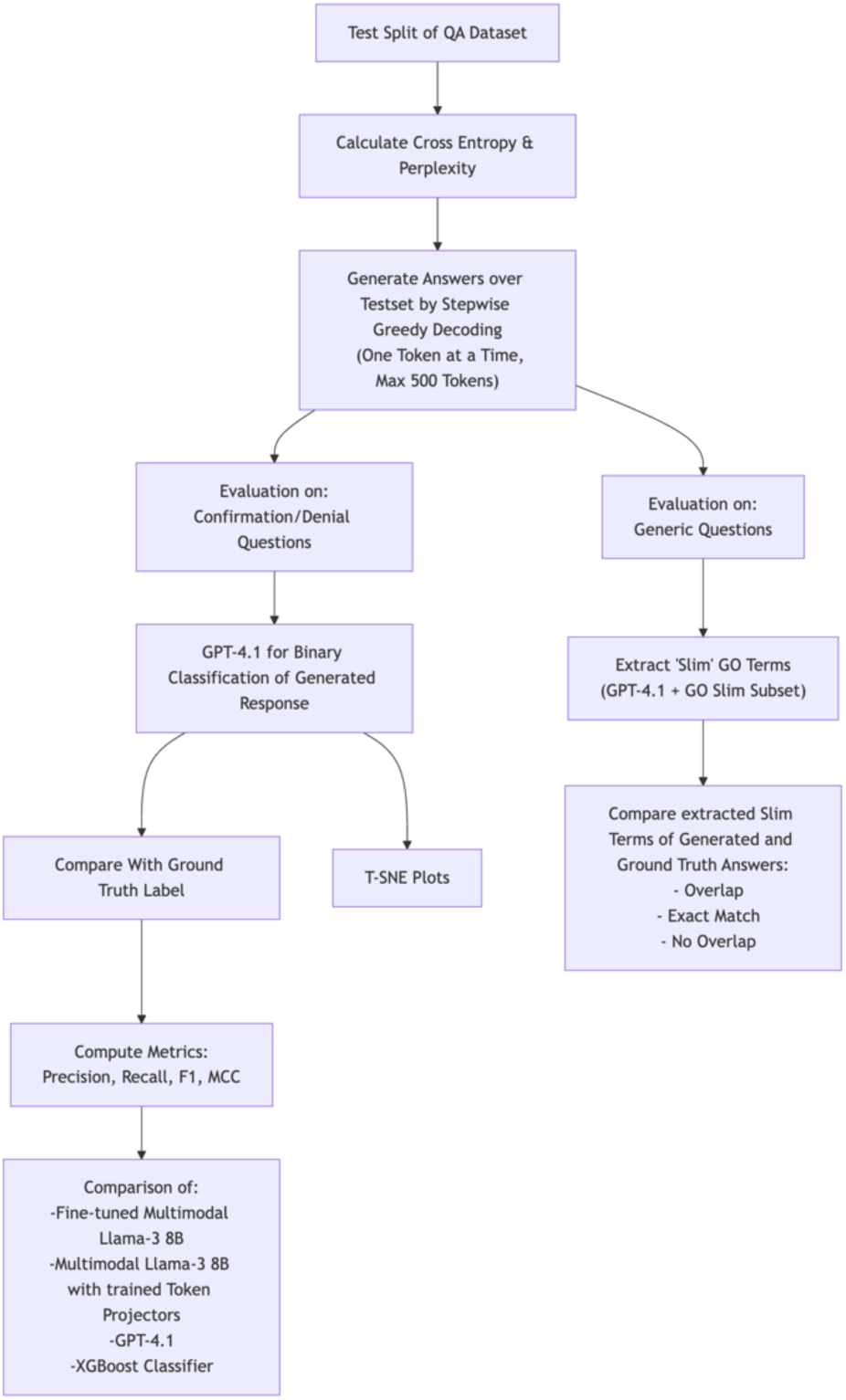
Model Evaluation Workflow using the test split of the QA Dataset

In addition to this joint fine-tuning approach, we also conducted a comparative experimental setup in which the Llama^33^ backbone was kept fixed and only the token projectors were trained. This enabled us to assess the impact of updating the core language model versus solely adapting the modality-specific projectors.

### Similarity-Aware Dataset Splitting

Initially, we adopted a conventional gene-based split, ensuring that the gene names assigned to the training, validation, and test sets were mutually exclusive. This strict partitioning guarantees that information pertaining to a given gene is confined to a single set, thus precluding direct data leakage and overestimation of model performance due to gene overlap.

However, restricting the split by gene names alone may not suffice to fully obviate information leakage, as homologous genes or genes with similar structural or sequence features may persist across splits. This realization motivated the adoption of a more robust, similarity-aware partitioning approach described below:

To prevent data leakage not just at the gene level but also across related sequences and structures, we adopted a similarity-aware partitioning protocol based on clustering in the multimodal embedding space. For each gene, the DNA, protein, and structure embeddings were concatenated to form a unified multimodal representation. These vectors were normalized and clustered via KMeans^38^ (k=3) using cosine similarity to ensure that structurally or functionally similar genes would be assigned to the same cluster.

Clusters were then mapped to train, validation, and test splits, with cluster sizes adjusted if necessary to achieve an approximate 80%/10%/10% data division. Assignment to each split was further refined using distance to cluster centroids, prioritizing samples most central to each cluster and ensuring exclusivity of samples across splits. This approach reduces the risk of intergenic data leakage, yielding more robust evaluations of model generalizability.

### Fine-Tuning and Experimental Setup

For efficiency, the training split was divided into three further equal-sized subsets. During each epoch, two of these were used, exposing every training point at least once every three epochs. Training was conducted for up to ten epochs, with a batch size of eight, a learning rate of 5 ∗ 10^−5^ and employed early stopping (patience = two epochs) based on validation loss. Model validation was performed after each epoch, with final selection and reporting based on results from the strictly held-out test set. The training was conducted using PyTorch^60^ and MLFlow^61^ was used for experiment tracking.

### Model Evaluation

#### A) Confirmation/Denial Task (Binary Classification)

To assess each model’s ability to confirm or deny an association between a given (unknown) sequence and a specified biological function, process, or component, we framed evaluation as a binary classification task. The following four models were compared:

- Multimodal-Llama (Full Fine-Tuning): Our multimodal Llama model in which both the virtual token projectors and the Llama backbone were fine-tuned.
- Multimodal-Llama (Projector-Only Fine-Tuning): An ablated variant where only the token projectors were trainable, and the Llama backbone was held fixed.
- XGBoost Binary Classifier: A non-generative baseline leveraging concatenated prompt, DNA, amino acid, and structure embeddings as input features for standard binary classification.
- GPT-4.1 Prompted Classifier: A system-prompted GPT-4.1 model tasked with direct classification using the DNA and amino acid sequences appended as plain text in the prompt.

### Evaluation Protocol

- Each model was prompted using question-answer test samples from the held-out KMeans split.
- For generative models, responses were labeled as “confirmation” or “denial” by an automatic GPT-4.1 post-processor.
- Predictions from all models were compared to ground truth binary labels.
- Performance was assessed using precision, recall, F1-score, and Matthews correlation coevicient (MCC).

#### B) Reasoning Assessment on Generic QA (Open-Ended Questions)

To evaluate the capacity for generative reasoning, only the fine-tuned multimodal Llama variants (with and without Llama adaptation) were considered. Here, models were prompted with open-ended questions regarding the possible biological functions, processes, or components associated with specific sequences.

##### Evaluation Approach

Generated responses were analyzed using GPT-4.1 and a Gene Ontology (GO) Slim subset, extracting a simplified set of GO terms from both model and reference answers.

### Overlap and accuracy were evaluated using

- The fraction of samples with overlapping model-generated and reference GO Slim terms.
- The fraction with exact set matches.
- The fraction with no overlap.

This two-pronged evaluation strategy provided both quantitative benchmarking of classification performance (across all models) and qualitative assessment of the generative reasoning capabilities of multimodal Llama models.

### Model Interpretability and Representation Analysis

To gain deeper insights into the internal mechanisms and learned representations of our models, we employed two complementary interpretability analyses: visualization of hidden states via t-distributed stochastic neighbour embedding (t-SNE)^43^ and a fine-grained examination of attention weights across modalities.

### Feature Space Visualization with t-SNE

To assess whether the model effectively integrated natural language and biological information, we extracted the hidden representations from the final Llama layer for each input-response pair. The mean or aggregated hidden states for each sequence were projected into two-dimensional space via t-SNE, allowing us to visualize the structure and clustering of learned embeddings. This analysis aimed to reveal whether the model developed a feature space in which natural language and biological contextual information were meaningfully combined, and whether generalizable genomic or functional rules emerged from fine-tuning.

### Model Interpretability via Attention Weights

To elucidate the model’s internal reasoning and to assess the influence of each modality during answer generation, we further performed a detailed analysis of attention weights focused on the virtual token representations. During inference on the test set, we captured the attention weights from the last hidden layer of the model at each generation step. Thus, we computed the average attention devoted to each modality within a response. This provided a summary measure or hint of which sources the model relied on most strongly during the generation. By quantifying attention patterns in this manner, our interpretability framework enabled us to characterize not only the integration of multimodal knowledge, but also the contributions of DNA, amino acid sequence, and structure information in guiding the model’s answer generation.

## Abbreviations

LLMs: Large Language Models
QA: Question-Answer
DNA: Deoxyribonucleic Acid
GO: Gene Ontology
GPT: Generative Pretrained Transformer
Evo2: DNA Foundation Model
ESM: Protein Language Model
MCC: Matthews Correlation Coefficient
TP: Token Projector
XGBoost: Extreme Gradient Boosting
BioBERT: Biomedical Bidirectional Encoder Representations from Transformers
GPU: Graphics Processing Unit
BRCA1: Breast Cancer Type 1 Susceptibility Protein
IHH: Indian Hedgehog Protein
INS: Insulin
BLAST: Basic Local Alignment Search Tool
t-SNE: t-distributed Stochastic Neighbor Embedding
MANE: Matched Annotation from NCBI and EMBL-EBI
PDB: Protein Data Bank
GCAE: Graph Convolutional Autoencoder

## 7. Declarations

### Ethics approval and consent to participate

Not applicable

## Consent for publication

Not applicable

## Availability of data and materials

The code developed for this study will be made publicly available on GitHub upon publication. The datasets generated and used for training the models, as well as the trained model, will be released on the Hugging Face platform. All resources will be accessible to the research community for reproducibility and further research.

## References

1. Lowe, J. B. & Marth, J. D. A genetic approach to Mammalian glycan function. Annu. Rev. Biochem. 72, 643–691 (2003).

2. Wright, B. W., Yi, Z., Weissman, J. S. & Chen, J. The dark proteome: translation from noncanonical open reading frames. Trends Cell Biol. 32, 243–258 (2022).

3. Callaway, E. ‘Dark proteins’ hiding in our cells could hold clues to cancer and other diseases. Nature 637, 1038–1040 (2025).

4. Reid Cahn, A., Bhardwaj, N. & Vabret, N. Dark genome, bright ideas: Recent approaches to harness transposable elements in immunotherapies. Cancer Cell 40, 792–797 (2022).

5. Blanco-Melo, D. et al. A novel approach to exploring the dark genome and its application to mapping of the vertebrate virus fossil record. Genome Biol. 25, 120 (2024).

6. Kreitz, J. et al. Programmable protein delivery with a bacterial contractile injection system. Nature 616, 357–364 (2023).

7. Kulmanov, M. et al. Protein function prediction as approximate semantic entailment. Nat. Mach. Intell. 6, 220–228 (2024).

8. Wang, B. et al. ProtGO: universal protein function prediction utilizing multi-modal gene ontology knowledge. Bioinformatics 41, btaf390 (2025).

9. Boadu, F. & Cheng, J. Improving protein function prediction by learning and integrating representations of protein sequences and function labels. Bioinforma. Adv. 4, vbae120 (2024).

10. Gligorijević, V. et al. Structure-based protein function prediction using graph convolutional networks. Nat. Commun. 12, 3168 (2021).

11. Wang, H., Zheng, H. & Chen, D. Z. TANGO: A GO-Term Embedding Based Method for Protein Semantic Similarity Prediction. IEEE/ACM Trans. Comput. Biol. Bioinform. 20, 694–706 (2023).

12. Zhong, X., Kaalia, R. & Rajapakse, J. C. GO2Vec: transforming GO terms and proteins to vector representations via graph embeddings. BMC Genomics 20, 918 (2019).

13. Merino, G. A., Saidi, R., Milone, D. H., Stegmayer, G. & Martin, M. J. Hierarchical deep learning for predicting GO annotations by integrating protein knowledge. Bioinformatics 38, 4488–4496 (2022).

14. Zhao, Y., Fu, G., Wang, J., Guo, M. & Yu, G. Gene function prediction based on Gene Ontology Hierarchy Preserving Hashing. Genomics 111, 334–342 (2019).

15. Wang, W. et al. A comprehensive computational benchmark for evaluating deep learning-based protein function prediction approaches. Brief. Bioinform. 25, bbae050 (2024).

16. Ieremie, I., Ewing, R. M. & Niranjan, M. TransformerGO: predicting protein–protein interactions by modelling the attention between sets of gene ontology terms. Bioinformatics 38, 2269–2277 (2022).

17. Raza, M., Jahangir, Z., Riaz, M. B., Saeed, M. J. & Sattar, M. A. Industrial applications of large language models. Sci. Rep. 15, 13755 (2025).

18. Jin, Q., Yang, Y., Chen, Q. & Lu, Z. GeneGPT: augmenting large language models with domain tools for improved access to biomedical information. Bioinformatics 40, btae075 (2024).

19. Wang, S., et al. Large Language Models for Education: A Survey and Outlook. Preprint at 10.48550/arXiv.2403.18105 (2024).

20. Masri, S., Ashqar, H. I. & Elhenawy, M. Leveraging Large Language Models (LLMs) for Traffic Management at Urban Intersections: The Case of Mixed Traffic Scenarios. Preprint at 10.48550/arXiv.2408.00948 (2024).

21. Li, Y., Wang, S., Ding, H. & Chen, H. Large Language Models in Finance: A Survey. Preprint at 10.48550/arXiv.2311.10723 (2024).

22. Brixi, G. et al. Genome modeling and design across all domains of life with Evo 2. 2025.02.18.638918 Preprint at 10.1101/2025.02.18.638918 (2025).

23. Nguyen, E. et al. Sequence modeling and design from molecular to genome scale with Evo. Science 386, eado9336 (2024).

24. Dalla-Torre, H. et al. Nucleotide Transformer: building and evaluating robust foundation models for human genomics. Nat. Methods 22, 287–297 (2025).

25. Sanabria, M., Hirsch, J., Joubert, P. M. & Poetsch, A. R. DNA language model GROVER learns sequence context in the human genome. Nat. Mach. Intell. 6, 911–923 (2024).

26. Nguyen, E., et al. HyenaDNA: Long-Range Genomic Sequence Modeling at Single Nucleotide Resolution. Preprint at 10.48550/arXiv.2306.15794 (2023).

27. J, Z. et al. Cross-species modeling of plant genomes at single-nucleotide resolution using a pretrained DNA language model. Proc. Natl. Acad. Sci. U. S. A. 122, (2025).

28. de Almeida, B. P. et al. A multimodal conversational agent for DNA, RNA and protein tasks. Nat. Mach. Intell. 7, 928–941 (2025).

29. Ashburner, M. et al. Gene Ontology: tool for the unification of biology. Nat. Genet. 25, 25–29 (2000).

30. Danner, M., Begemann, M., Elbracht, M., Kurth, I. & Krause, J. Utilizing protein structure graph embeddings to predict the pathogenicity of missense variants. 2024.11.15.623748 Preprint at 10.1101/2024.11.15.623748 (2024).

31. Guarrasi, V. et al. A systematic review of intermediate fusion in multimodal deep learning for biomedical applications. Image Vis. Comput. 158, 105509 (2025).

32. Wang, J., et al. A Comprehensive Review of Multimodal Large Language Models: Performance and Challenges Across Different Tasks. arXiv.org https://arxiv.org/abs/2408.01319v1 (2024).

33. Grattafiori, A., et al. The Llama 3 Herd of Models. Preprint at 10.48550/arXiv.2407.21783 (2024).

34. Touvron, H., et al. Llama 2: Open Foundation and Fine-Tuned Chat Models. arXiv.org https://arxiv.org/abs/2307.09288v2 (2023).

35. Touvron, H., et al. LLaMA: Open and Efficient Foundation Language Models. arXiv.org https://arxiv.org/abs/2302.13971v1 (2023).

36. Lin, Z. et al. Evolutionary-scale prediction of atomic-level protein structure with a language model. Science 379, 1123–1130 (2023).

37. OpenAI, et al. GPT-4 Technical Report. Preprint at 10.48550/arXiv.2303.08774 (2024).

38. Ikotun, A. M., Ezugwu, A. E., Abualigah, L., Abuhaija, B. & Heming, J. K-means clustering algorithms: A comprehensive review, variants analysis, and advances in the era of big data. Inf. Sci. 622, 178–210 (2023).

39. Chicco, D. & Jurman, G. The Matthews correlation coefficient (MCC) should replace the ROC AUC as the standard metric for assessing binary classification. BioData Min. 16, 4 (2023).

40. Chen, T. & Guestrin, C. >XGBoost: A Scalable Tree Boosting System. Preprint at 10.48550/arXiv.1603.02754 (2016).

41. Lee, J. et al. BioBERT: a pre-trained biomedical language representation model for biomedical text mining. Bioinformatics 36, 1234–1240 (2020).

42. Sf, A., W, G., W, M., Ew, M. & Dj, L. Basic local alignment search tool. J. Mol. Biol. 215, (1990).

43. Maaten, L. van der & Hinton, G. Visualizing Data using t-SNE. J. Mach. Learn. Res. 9, 2579–2605 (2008).

44. Wang, S., Hu, M., Li, Q., Safari, M. & Yang, X. Capabilities of GPT-5 on Multimodal Medical Reasoning. arXiv.org https://arxiv.org/abs/2508.08224v2 (2025).

45. Leon, M. GPT-5 and open-weight large language models: Advances in reasoning, transparency, and control. Inf. Syst. 136, 102620 (2026).

46. Team, G., et al. Gemini: A Family of Highly Capable Multimodal Models. Preprint at 10.48550/arXiv.2312.11805 (2025).

47. Comanici, G., et al. Gemini 2.5: Pushing the Frontier with Advanced Reasoning, Multimodality, Long Context, and Next Generation Agentic Capabilities. Preprint at 10.48550/arXiv.2507.06261 (2025).

48. Hinton, G., Vinyals, O. & Dean, J. Distilling the Knowledge in a Neural Network. Preprint at 10.48550/arXiv.1503.02531 (2015).

49. Cooper, N. & Scholak, T. Perplexed: Understanding When Large Language Models are Confused. arXiv.org https://arxiv.org/abs/2404.06634v1 (2024).

50. Muhlgay, D. Generating Benchmarks for Factuality Evaluation of Language Models.

51. Dathathri, S., et al. Plug and Play Language Models: A Simple Approach to Controlled Text Generation. Preprint at 10.48550/arXiv.1912.02164 (2020).

52. Liu, C.-W. et al. How NOT To Evaluate Your Dialogue System: An Empirical Study of Unsupervised Evaluation Metrics for Dialogue Response Generation. in Proceedings of the 2016 Conference on Empirical Methods in Natural Language Processing 2122–2132 (Association for Computational Linguistics, Austin, Texas, 2016). doi:10.18653/v1/D16-1230.

53. Morales, J. et al. A joint NCBI and EMBL-EBI transcript set for clinical genomics and research. Nature 604, 310–315 (2022).

54. Cock, P. J. A. et al. Biopython: freely available Python tools for computational molecular biology and bioinformatics. Bioinformatics 25, 1422–1423 (2009).

55. Jumper, J. et al. Highly accurate protein structure prediction with AlphaFold. Nature 596, 583–589 (2021).

56. Abramson, J. et al. Accurate structure prediction of biomolecular interactions with AlphaFold 3. Nature 630, 493–500 (2024).

57. Varadi, M. et al. AlphaFold Protein Structure Database in 2024: providing structure coverage for over 214 million protein sequences. Nucleic Acids Res. 52, D368–D375 (2023).

58. Wolf, T., et al. HuggingFace’s Transformers: State-of-the-art Natural Language Processing. Preprint at 10.48550/arXiv.1910.03771 (2020).

59. Jamasb, A. et al. Graphein - a Python Library for Geometric Deep Learning and Network Analysis on Biomolecular Structures and Interaction Networks. Adv. Neural Inf. Process. Syst. 35, 27153–27167 (2022).

60. Paszke, A., et al. PyTorch: An Imperative Style, High-Performance Deep Learning Library. Preprint at 10.48550/arXiv.1912.01703 (2019).

61. Chen, A. et al. Developments in MLflow: A System to Accelerate the Machine Learning Lifecycle. in Proceedings of the Fourth International Workshop on Data Management for End-to-End Machine Learning 1–4 (Association for Computing Machinery, New York, NY, USA, 2020). doi:10.1145/3399579.3399867.

